# Growth charts of infant visual neurodevelopment generalize across global contexts

**DOI:** 10.1101/2025.03.25.645314

**Authors:** Emma T. Margolis, Chris Camp, Ana C. Sobrino, Guilherme V. Polanczyk, Daniel Fatori, Khula South Africa Team, Germina Team, LABS Team, GABA Team, 1kD EEG Working Group, 1kD Machine Learning Working Group, Laura Cornelissen, Charles B. Berde, Takao K. Hensch, Charles A. Nelson, Elizabeth Shephard, Kirsten A. Donald, Dustin Scheinost, Laurel J. Gabard-Durnam

**Author notes:** **Corresponding Authors:** Emma Margolis,; Laurel Gabard-Durnam. indicates equal contributions (Emma T. Margolis, Chris Camp). indicates equal contributions (Dustin Scheinost, Laurel J. Gabard-Durnam).

## Abstract

Normative brain growth charts in early life hold great promise for furthering basic and clinical science. We leverage the rapid, substantial development of visual cortex function that is indexed by visual-evoked potentials (VEP) in electroencephalography to create longitudinal normative growth curves of task-related brain function with 1374 observations contributed by 802 infants (57 to 579 days old) from South Africa, Brazil, and the United States. Site-specific models were cross-validated and showed excellent fits to other sites’ samples, demonstrating functional growth curves generalize across contexts robustly. Deviations from the normative growth models associated with early environmental and behavioral measures such as prenatal exposures and postnatal cognition. These findings demonstrate the utility of using functional growth charts to understand and potentially act on individual neurodevelopmental trajectories. VEP brain function growth charts represent a new direction for EEG research to support healthy brain development globally.

## Introduction

Normative changes quantified through physical growth charts have transformed science, public health, and pediatric care^1–5^. The brain undergoes tremendous functional and structural change across the lifespan^6–12^. Normative brain growth charts also hold great promise for furthering basic and clinical science^13–17^. Growth charts may reveal milestones in brain development and provide individual-level neural indicators facilitating risk identification, prognosis, stratification, or intervention evaluation^13,14,18,19^. Several candidate growth charts for normative brain structure^13,18,20^ and function at rest or in sleep are emerging for the lifespan^21–23^. Growth charts characterizing awake brain function during task paradigms can offer unique information about how brains function and change. This approach may also improve chart utility as task-related activity, compared to resting activity, has shown greater sensitivity and specificity in associating individual differences in brain function with cognition, symptoms, and behavior^24,25^. However, normative growth charts of task-related brain function have yet to be constructed.

There are multiple challenges in identifying which charts of task-related brain function are feasible and useful^13,17^. These include identifying standardized, scalable, and geoculturally-appropriate paradigms to elicit specific neural activity across contexts and development^26–33^. Statistical strategies must also facilitate external validation of growth curves to test their robustness across contexts and integrate information from multiple features generated during the task^17^. Finally, while initial brain growth charts have taken a lifespan approach, growth curves have poorer performance at their tails, meaning brain changes in infancy may be less well-characterized^34^. However, elevated early neuroplasticity enables rapid, substantial brain development^35–38^. As mandated for physical growth charts, targeted brain charts with dense longitudinal sampling in early life can best capture these changes^4,39^.

Here, we address these challenges to construct and evaluate brain function growth charts in awake infants. We focus on characterizing the rapid, early development of visual cortical function that can be precisely and efficiently measured through a standardized phase-reversal visual-evoked potential (VEP) paradigm across contexts and ages^29–32,40–43^. Charting visual neurodevelopment provides critical information about foundational sensory development. It also provides early functional indicators of risk, scaffolding for subsequent higher-order cognition, and opportunities for translational research across species^37,43–46^. We chart longitudinal data (1374 observations) over the first two years of life from four cohorts across three continents that represent distinct cultural, linguistic, geographic, and socioeconomic contexts (Cape Town, South Africa, São Paulo, Brazil, and Boston, USA) with *a priori* harmonization of paradigm, data acquisition, and processing parameters^47–49^. We leverage electroencephalography (EEG) as an ideal method to capture standardized features of awake brain function in early life^26,28,29^ and generalized additive models of location, scale, and space (GAMLSS) with external cross-validation to create and generalize growth curves^4,17,50^.We demonstrate that brain function growth charts in infancy are feasible, stable, and generalizable. We also show that they capture individual deviations that are both sensitive to early developmental risk factors and relate to emerging behavior.

## Results

We constructed developmental growth curves of visual brain function from 57 to 579 days of age with participants (n=802) from four longitudinal cohorts (i.e., Khula South Africa, Germina, LABS, GABA) across three continents (i.e., Africa, South America, and North America) (Fig. 1). Cohorts’ characteristics are summarized in Table 1, and demographic information for each cohort is summarized in Table S1. Paradigm, acquisition, and processing were harmonized *a priori* across all sites, and processing parameters are available in Tables S2 and S3. Visual brain function was indexed by 128-channel electroencephalography (EEG)- derived pattern-reversal visual-evoked potentials (VEP) using a standard black-white checkerboard paradigm on an external monitor while infants sat on their caregiver’s lap^29,51^. VEPs were extracted and quantified from an occipital ROI (Fig. 2B) over the primary visual cortex generator^52–57^ with respect to six standard morphological features—peak amplitudes and latencies to those peak amplitudes (i.e., latencies) for the N1, P1, and N2 components (Fig. 2A)^58^. These morphological features reflect distinct functional and structural processes that characterize primary visual cortical development. Specifically, N1 and N2 components reflect parvocellular visual pathway development (sensitive to color and spatial detail), while the P1 component is generated by the magnocellular pathway (sensitive to motion)^59–64^. Amplitude features reflect cortical neurotransmission, especially the balance of excitatory/inhibitory postsynaptic potentials^58,65–67^, while latencies largely reflect the structural integrity and myelination of the visual pathways^51,68,69^. We modeled the six VEP features independently using generalized additive models of location, scale, and space (GAMLSS), a World Health Organization-recommended framework to model non-linear growth trajectories^4^.

**Figure 1:**
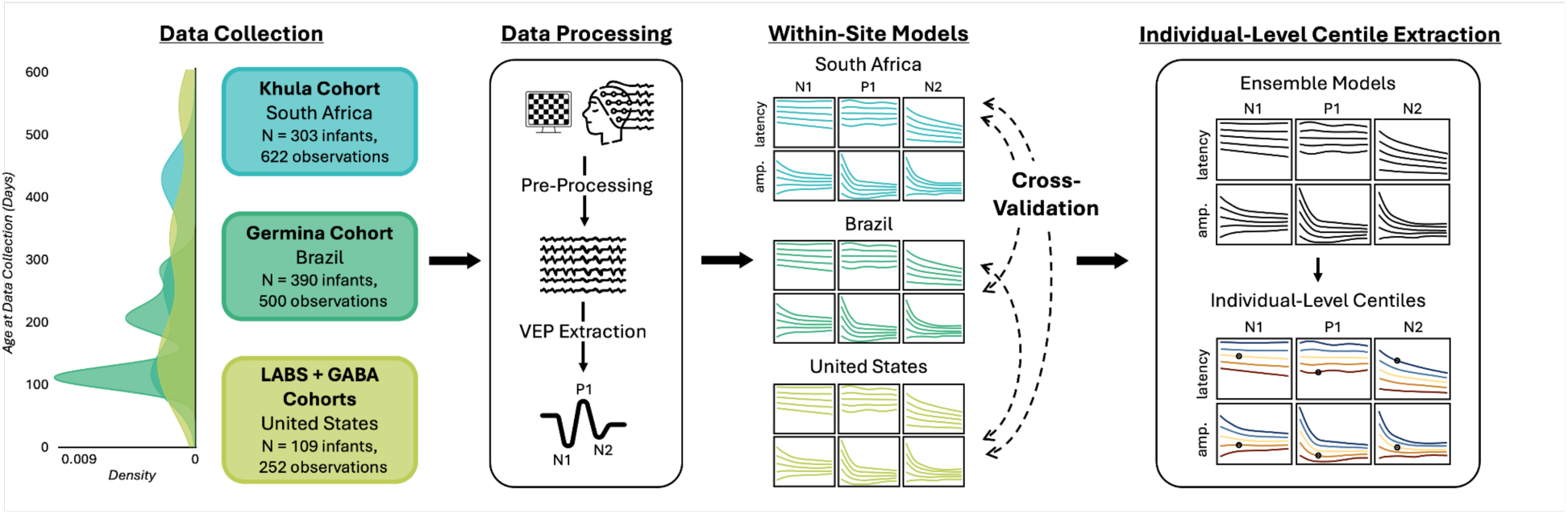
Study Overview. We created longitudinal growth curves using data from 802 infants (*n*=1374 EEG observations) between 57 and 579 days old across three continents. 128-channel EEG data were *a priori* harmonized for paradigm, acquisition, and processing parameters across sites. We independently trained GAMLSS curves for each site and VEP feature—amplitudes and latencies for the N1, P1, and N2 components. The GAMLSS curves were externally cross-validated across sites. Individual centiles were extracted and associated with a known developmental risk factor (prenatal alcohol exposure) and subsequent cognitive scores on a global public health measure (Global Scales of Early Development, Long Form).

**Figure 2:**
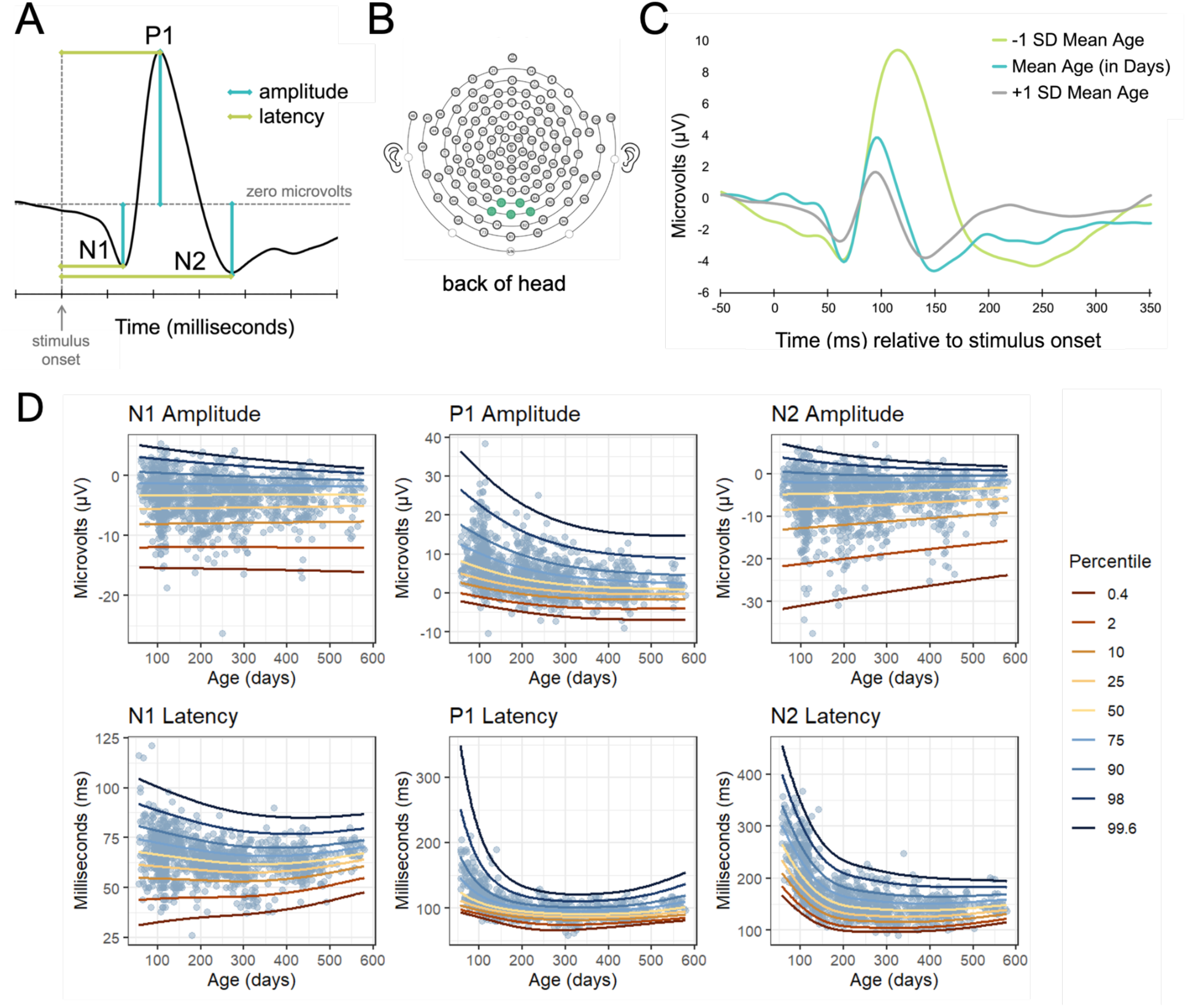
Characterizing the VEP Over Infancy. (A) Demonstrative VEP waveform with labeled components (i.e., N1, P1, N2) and features (i.e., amplitude, latency). (B) 128-channel electrode configuration with highlighted occipital region of interest (i.e., E70, E71, E75, E76, E83). (C) Averaged VEP waveforms across infants (n=15 each) for the mean age of our dataset (in days) and ± 1 SD (i.e., 227.66, 353.23, 102.09 days). (D) Growth curves for VEP features trained on all three sites (*n* = 1371 observations). Nonlinear changes are primarily observed in the P1 amplitude and the P1 and N2 latencies.

**Table 1:** Cohort Characteristics.

| Cohort | Site | Age Range (Days) | N infants | N infants Providing >1 Useable Timepoint | N observations |
| --- | --- | --- | --- | --- | --- |
| Khula | Cape Town, South Africa | [58, 516] | 303 | 214 (70.63%) | 622 |
| Germina | São Paulo, Brazil | [94, 312] | 390 | 110 (28.21%) | 500 |
| LABS | Boston, USA | [85, 579] | 56 | 38 (67.86%) | 157 |
| GABA | Boston, USA | [57, 366] | 53 | 33 (62.26%) | 95 |

### Generalizing Growth Curves of Visual Neurodevelopment

We trained GAMLSS curves for each VEP feature to model normative development of visual cortex function across sites. The fitting and testing procedures are outlined in Fig. S1. Models were fit with p-spline smoothing terms and fixed effects for each feature. Amplitudes were shifted by their minimum to have the range [1, ∞). We prioritized developing models generalizable across all three sites. Distributions for each VEP feature model were chosen by minimizing the Bayesian information criterion (BIC) in the held-out site of a cross-validation fold. Smoothing penalties were tuned by optimizing the generalized Akaike information criterion penalty *k* to minimize the total validation BIC. Using the established hyperparameters, we fit models on all pairs of site datasets. Models were assessed by examining the residuals’ worm plots and *Q* statistics and comparing the predicted and actual VEP feature values with Pearson *r* correlation and root mean square error (RMSE). 95% confidence intervals were generated for correlations with 1000-fold nonparametric bootstraps.

All VEP feature curves exhibited high model fits, with an average RMSE of 1.13 *μ*V for amplitudes and 13.24 ms for latencies. Similarly, the average Pearson correlation across site datasets between the predicted and observed z-scores ranged between *r=*.68 (N2 Latency, 95% CI: .64–.72) to *r*=1 (N1 Amplitude, 95% CI: .99–1). Additional model statistics are in Table S5. We did not observe significant differences in correlations across site pairs with an omnibus ANOVA (*F*(2)=.15, *p*=.86), suggesting that the models fit site pairs equally well. Amplitudes had higher correlations between models and data than latencies (two-sided *t* test: *t*(14.36)=2.95, *p*<.05), likely due to their simpler distributions. Including sex as a covariate did not change model fit for any of the VEP features. Therefore, it was not included in any of the models.

#### Models Generalize Across Sites

To assess generalizability across sites, we then evaluated model fit in the held-out dataset for each site pair. All feature curves exhibited high model fits on the held-out datasets, with an average RMSE of 2.16 *μ*V for amplitudes and 15.94 ms for latencies (Table S5). The average Pearson correlation across datasets between the predicted and observed z-scores ranged between *r=*.71 (N2 Latency, 95% CI: .66–.76) to *r*=.99 (N1 Amplitude, 95% CI: .99–1). As with the within-dataset fits, we did not observe significant differences in correlations across held-out datasets with an omnibus ANOVA (*F*(2)=.21, *p*=.81).

### Characterizing VEP Development

Next, we characterized the shape of the VEP growth curves to identify developmental trends in visual function across individuals. We trained a model for each feature on all available data to provide the best-powered representation of the growth curves (Table S6-7). The six VEP features exhibited different growth shapes, representing distinct developmental patterns in their trajectories (Fig. 2D). The best underlying distributions as determined by cross-validation were as follows: sinh-arcsinh (SHASH) for N1 amplitude; Johnson’s S_U_ for P1 amplitude, N2 amplitude, and P1 latency; *t*-family (TF) for N1 latency, and generalized inverse Gaussian (GIG) for N2 Latency. We plotted the fitted terms to examine the behavior of each parameter for the models and used numerical derivation to determine slope minima and maxima (Table S8, Fig. S14-23). We observed several trends across the features. The central tendency (*μ*, the ‘location’ parameter) of the amplitude peaks diminished with age, with the negative component amplitudes (N1, N2) increasing and P1 decreasing. Negative component amplitudes (N1, N2) demonstrated steeper slopes of change than the P1 component during infancy. Latency central tendency parameters also decreased with age, with N2 showing the steepest slopes over infancy. Variance (*σ*, the ‘scale’ parameter) decreased with age in all features except P1 latency as the components matured and converged. The remaining two parameters, *v* and *τ*, were more variable across features as they represent different moments depending on the distribution across individuals.

We also examined differences in VEP development to identify milestones in early visual function (Fig. 2C, S24-25). We estimated the 50th percentile of each feature across the age range and calculated the first and second derivatives (Fig. S26-29). As is expected, we observed decreasing latencies over development, indicating increased processing speed and myelination as infants get older^58^. P1 and N2 latencies had nonlinear shapes, with P1 latency maturing faster. We observed minimal change in N1 latencies over infancy. The magnitudes of amplitudes decreased with age, with negative amplitudes (i.e., N1 and N2) becoming less negative and the positive amplitude (i.e., P1) becoming less positive. P1 amplitude exhibited a nonlinear shape with the greatest changes in the first months of infancy. In contrast, the rates of change in N1 and N2 amplitudes (decreasing) were relatively constant over infancy, suggesting protracted change.

### An Individual’s Curve is Stable Across Development

We leveraged the longitudinal study design to characterize the stability of each individual’s growth centiles. For a normative growth curve, most individuals should stay near or within their growth centile across timepoints. Growth curve stability is frequently quantified by binning centile lines at the 5th, 10th, 25th, 50th, 75th, 90th, and 95th percentiles, since volatility is attenuated at the fringes of the growth curves. Centiles for all VEP features were generally stable, with 75% of participants shifting no more than one centile line across the measured period (Fig. 3). The mean centile change per timepoint across all features was .75 lines (*SD*=0.62), or 14.45 centiles (*SD*=10.68). We observed no significant differences in stability across VEP features (*F*(5, 1272)=.87, *p*=.50), suggesting that features were consistent in their stability. We explored centile change as a function of age by fitting a locally estimated scatterplot smoothing (LOESS) curve to the relationship between age and centile change per day (Fig. S29). The point of greatest change was at the earliest age for all features. There was a consistent decrease in volatility across the first year of development.

**Figure 3:**
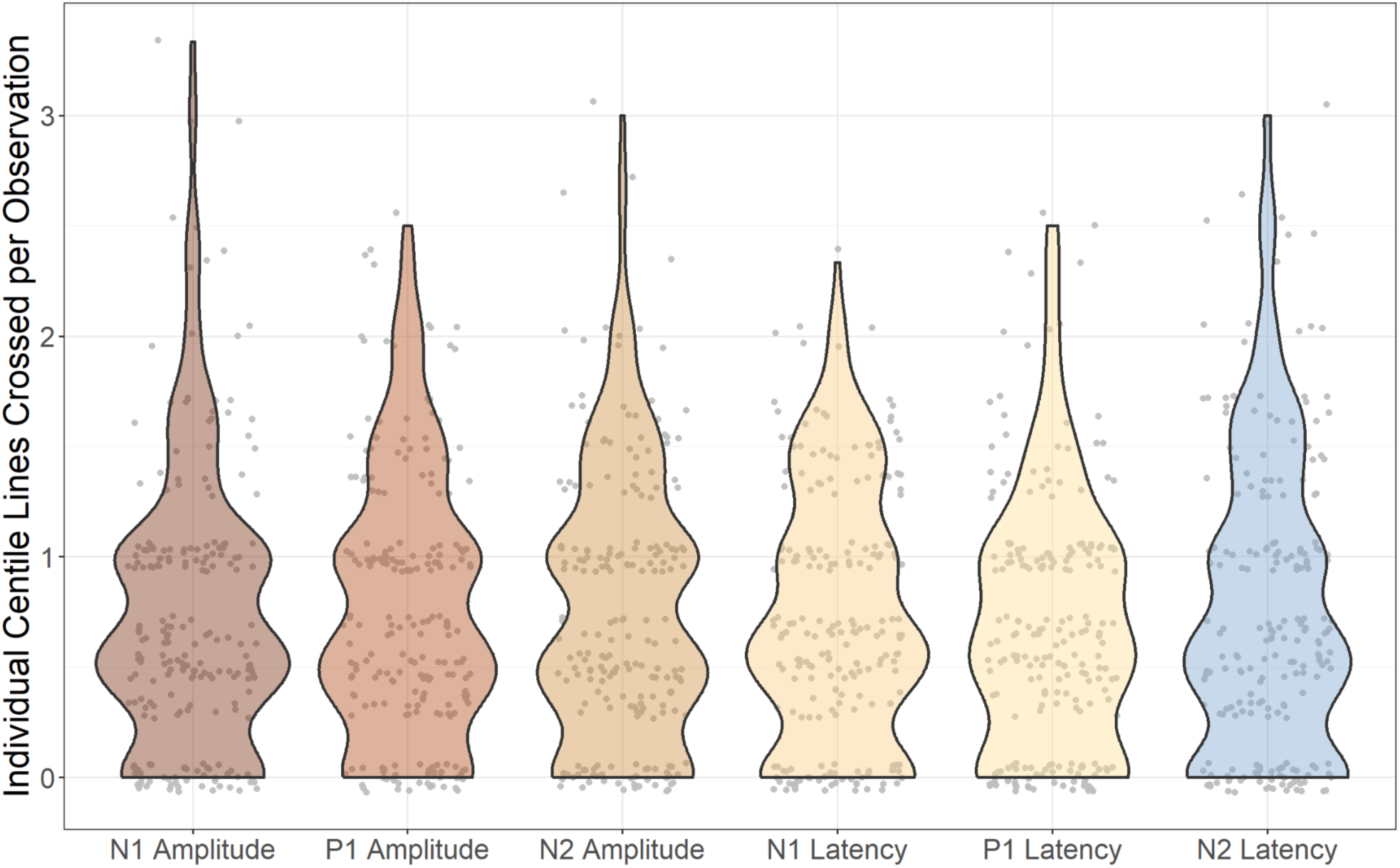
Stability of Individual VEP Centile Scores by Feature. Violin plot representing the number of centile lines (located at the 5th, 10th, 25th, 50th, 75th, 90th, and 95th percentiles) crossed by each individual per observation. 75% of participants shifted less than one centile line, indicating that most individuals remained near the same centile across infancy. No significant differences in stability across feature were observed.

### Assessing Utility of VEP Growth Charts: Associations with Cognitive Development and Early Exposures

#### Individual VEP Scores are Associated with Cognitive Development

We investigated whether individual differences in VEP development were associated with a globally scalable measure of neurodevelopment and emerging cognition designed for public health contexts, the Global Scales for Early Development (Long Form; GSED-LF), developed and validated by the WHO^70–72^. We tested whether deviation from the VEP GAMLSS curves related to the GSED-LF age-adjusted D (DAZ) scores measured at 14 months (mean age=429±30.66 days) in the South Africa site. As the VEP waveform includes six distinct features, each with a unique growth trajectory, we integrated the information contained within each feature into a composite deviation score per infant. We extracted z-scores for every observation and feature for each individual. These z-scores measure an individual’s deviation from the normative curve for that feature. We then summed all z-scores to derive an individual’s total deviation score. Using the models trained on the full dataset across all three sites, the VEP deviation scores were positively associated with GSED scores (*r*(89)=.27, *p*=.0089; Fig. 4A). This association remained when correcting for potential covariates of age at GSED collection and sex (*r*(89)=.22, *p*=.036), as well as percent VEP trials excluded during EEG quality control at each VEP observation (*r*(89)=.27, *p*=.012). To test the generalizability of this association, we applied the GAMLSS model trained in the Brazil and US sites to the held-out South Africa site to derive total deviation scores. Deviation scores were nearly identical to those derived from the model trained on all three sites (all *r*’s>.93, *p*<.001). Accordingly, the deviation scores were still associated with GSED scores (*r*(89)=.28, *p*=.0072). These results further demonstrate the ability of the VEP growth curves to generalize across populations.

**Figure 4:**
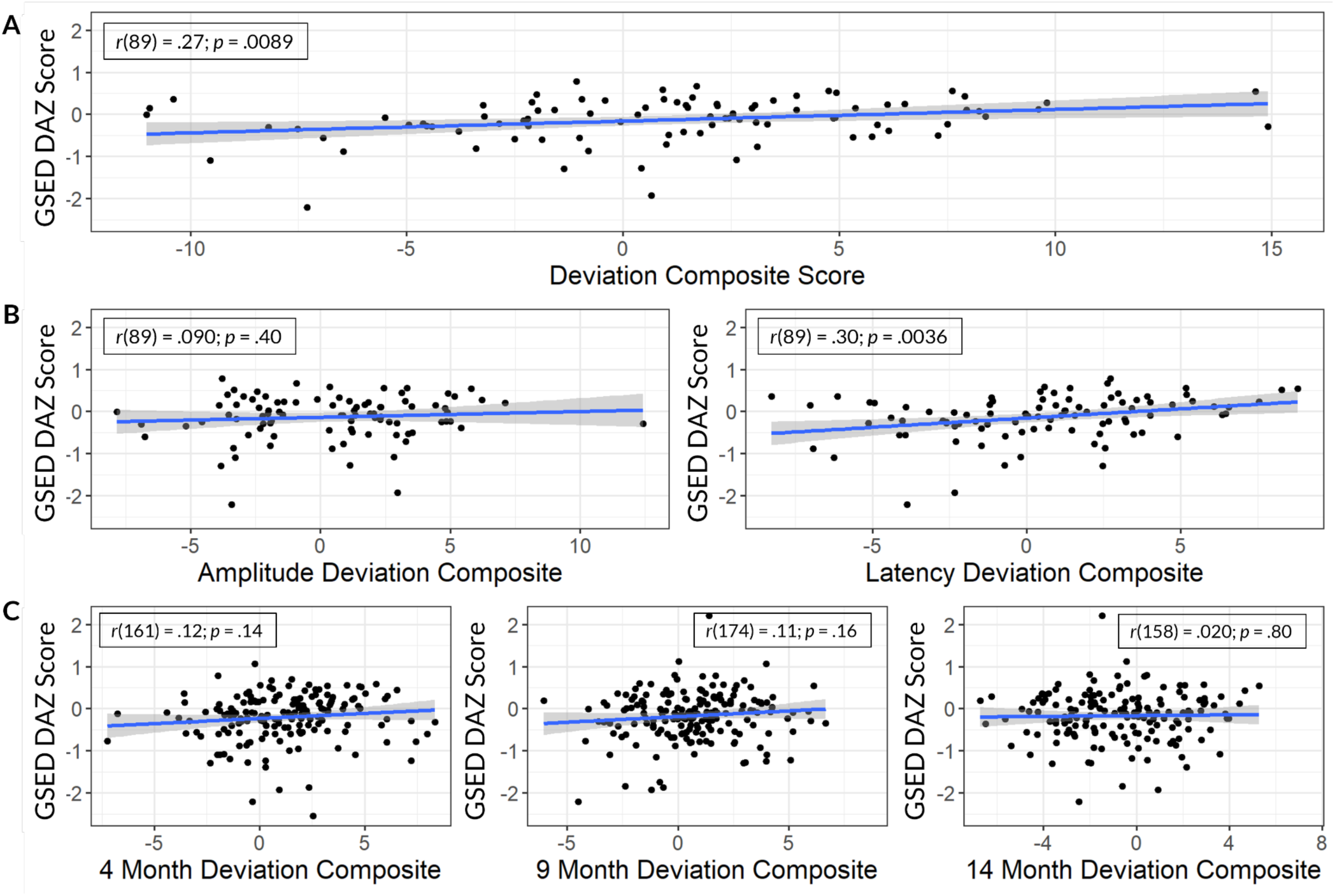
Associations of VEP Deviation scores and Cognitive Outcomes. Scatter plots representing associations between deviation composite scores and Global Scales for Early Development Long Form (GSED-LF) age-adjusted D (DAZ) scores measured at 14 months. A) Deviation scores across all features and measurements; B) Deviation composites separated by amplitude *vs*. latency; C) Deviations composites separated by age.

We conducted several *post hoc* analyses to better understand the association between growth curve deviations and neurocognitive development. We separated the total deviation scores by VEP latency and amplitude. Deviations from the latency feature curves were significantly associated with GSED scores (*r*(89)=.30; *p*=.0036), while amplitude deviations were not (*r*(89)=.090; *p*=.40), suggesting that the latency deviations were driving the association with GSED scores (Fig. 4B). We also separated the total deviation scores by age based on the target VEP observation timepoints. Deviations at a single age group were not associated with cognitive outcomes measured by the GSED (*r*’s<.12, *p’s*>.12), highlighting the improved power by integrating longitudinal data (Fig. 4C). This remained true when only using the latency measurements (*r*’s<.094, *p*’s>.23). Associations between all individual feature measurements and GSED scores are shown in Fig. S31-33.

#### Prenatal Alcohol Exposure is Associated with Deviations from Normative VEP Development

Finally, to test the sensitivity of VEP normative growth curve centiles to known environmental influences, we have replicated results from a prior study with the South African site that used conventional VEP analyses and a subset of the current sample. Specifically, we have previously shown that prenatal alcohol exposure during the first trimester was associated with longer P1 latencies in the first 6 months of life that neurally resolve by the end of the first year^51^. Here, we extracted P1 latency centile scores in the Khula South Africa dataset to replicate this finding. Corrected P1 latency centile scores for infants under 6 months positively associated with drinks per week during the first trimester (*r*(131)=.43, *p*<.001). Similarly to the original analysis, this effect was not observed at later ages (*r*(130)=-.022, *p*=.80). Results were identical when using the models with the South Africa site data held out of the training process (*r*(130)=.43, *p*<.001). This replication demonstrates the ability of centile curves to detect developmentally-sensitive effects of environmental influences, including those relevant to public health efforts, even in held-out site data.

## Discussion

We have created and validated brain function growth charts in awake infants over the first two years of life. In contrast to other extant brain charts^13,18,21^, we have used task activity and *a priori* harmonized, longitudinal data, demonstrating the feasibility of this approach to understand brain development at a global scale. In keeping with established practice for physical growth charts that single out limited, early periods of rapid growth^1–5^, we focused on the narrow but critical age range of the first two postnatal years to rigorously map the substantial brain changes underlying visual function during this period^58,73,74^. We targeted the visual-evoked potential (VEP) index of sensory brain function, which is neurobiologically richly-characterized, clinically relevant, and widely collected^29–32,40–43,51,51–57^. The growth charts generalized well across sites with distinct sociocultural and economic profiles from three continents. Deviations from the normative growth models associated with both environmental influences (prenatal alcohol exposure) and postnatal cognitive scores on a global WHO-validated public health measure, demonstrating growth charts’ utility to understand individual neurodevelopment and potential to integrate with early identification and intervention efforts. Overall, our brain function growth charts represent a new direction for EEG research and have broad impacts in the following ways.

### Generalized reference growth charts for early brain function

We prioritized scalability and generalizability across global contexts with diverse populations in this investigation so that our growth charts may serve as reference models for the broader community of neurodevelopmental researchers. Most of our participants live in global majority contexts (e.g., global South), which are underrepresented in neuroscience research^75–79^, including functional brain chart efforts^21,22^. Critically, the three sites have sufficient age range overlap and sample size to distinguish between site and age effects. Overall, this diversity provides a thorough test of the generalizability of our models. Other researchers can leverage these models to estimate centiles for each VEP feature from smaller cohorts, special populations, or after new data collection efforts without creating new growth charts. These centiles scores can be combined using our composite deviation score approach to increase power in downstream analyses. To facilitate these efforts, we integrated these growth charts directly into HAPPE—an open-source, user-friendly software for processing EEG data used in the present study^80,81^. We also make VEP stimuli, paradigm scripts, and processing parameter files available so researchers can replicate the VEP procedure and processing precisely and seamlessly progress within HAPPE from raw EEG data to growth-chart-derived centile scores for new data. The cost-effectiveness, tolerance, and scalability of EEG relative to other early life neuroimaging approaches further make this a promising approach to expand early neurodevelopmental research efforts.

### Insights into Early Visual Functional Neurodevelopment

The six VEP features showed divergent developmental patterns that invite future mechanistic work. The temporal pattern of changes differed between amplitude and latency features, with peak amplitudes exhibiting gentle slopes over infancy, while latencies show distinct nonlinear decreases. This pattern is consistent with separable underlying developmental mechanisms. Specifically, amplitude changes reflect postsynaptic dynamics in local circuits, including the balance of excitatory and inhibitory neurotransmission (E/I balance), which changes over infancy^58,65,66^. Synaptogenesis rates also increase in primary visual cortex during this infant window and impact local circuit dynamics^82,83^. VEP amplitude changes may provide functional indices of the interplay between rapid synaptogenesis and changing E/I balance within the primary visual cortex. In contrast, latency reductions reflect myelination of the geniculate-occipital pathways^68,69,84,85^, and MRI-derived myelin-sensitive measures have been related to infant VEP latency in a subsample of the study participants previously^51^. Here we show these latency changes largely resolve within the first year.

With respect to the different VEP components, the slopes of the N1 component peak amplitude and latency are largely flat within the featured age range, although the N1 latency exhibits a very gentle negative slope before postnatal day 200. This pattern suggests a very early and brief window of postnatal development for the N1 component within the parvocellular pathway^58^. In contrast, the P1 and N2 features showed significant developmental changes over the infant period, consistent with prior studies of visual function across imaging modalities^86^. Notably, the shallower slope of P1 amplitude change and earlier stabilization of the P1 latency relative to the N2 amplitude and latency patterns is consistent with earlier functional and structural maturation of the magnocellular relative to the parvocellular visual pathway in infancy as others have posited^60,61^. Further research is required to precisely link the underlying neurobiological mechanisms to observed developmental trajectories in these VEP growth charts. All features stabilized by 2 years, supporting our targeted focus on this developmental window (though protracted changes at smaller scales occur through childhood).

Additionally, we found empirical support for the role of early, basic visual neurodevelopment in scaffolding higher-level cognition. Though prior research has linked infant visual function behaviorally (e.g., visual attention) with cognition^87–89^, studies testing early visual brain function directly through the VEP have been inconclusive with respect to whether these early functional properties relate to cognition^41,90,91^. We found that deviations from normative VEP development were associated with a globally-validated public health index of cognition (the Global Scales of Early Development Long Form). This finding in the context of prior studies suggests that it is the trajectory of VEP development rather than discrete features or timepoints that relate to emerging cognition. Alterations in timing (i.e., precocious or delayed maturation) of the neurobiological processes underlying VEP development can have long-term consequences for behavior^45,92–96^. For example, alterations in the timing of sensory neurodevelopment are implicated in disorders such as Rett Syndrome^45,92,94,95,97^, Down Syndrome^92,95^, and Autism Spectrum Disorders^45,92,95^. Our results using normative modeling highlight the power of this approach to uncover early brain-behavior relations at the individual level during the infant period when early intervention support is most efficacious.

### Methodological Advances

Methodologically, we make several contributions to mapping normative growth charts in neuroimaging data. First, we borrowed modeling approaches from machine learning, excluding subsets of data (e.g., a site) during initial fitting to improve generalizability. Training and evaluating models in disparate populations prevents overfitting^98^. Our unique dataset with overlapping age ranges and *a priori* harmonization but distinct cultural, linguistic, geographic, and socioeconomic contexts facilitates this testing. Second, our longitudinal data allowed us to estimate centile stability by tracking changes within individuals following the existing literature. Our curves demonstrated better individual centile stability than other developmental growth curves, like weight^99^. In contrast, previous work with mainly cross-section neuroimaging data reported the linearly interpolated interquartile range, which discards individual level data by averaging^13^. Finally, we combined centiles from multiple timepoints and features to improve effect sizes with developmental outcomes. Simple multivariate approaches improve effect sizes beyond individual univariate features^100,101^. Notably, no single feature or timepoint was associated with cognitive outcomes on its own. Future work can assess how additional composite score calculations may further increase power across brain growth charts.

### Limitations

Unlike other brain growth charts, but in keeping with established physical growth charts like weight and height, we intentionally used a narrow age range (57–579 days) within the first two postnatal years. The visual cortex most rapidly changes structurally and functionally during this period^58,102,103^. Our results reinforce this premise, as all VEP features significantly slowed rates of change by the end of the second year of life. Nevertheless, changes in the VEP waveform also occur before 57 days and are not captured in our models. Our sample size (*n*=1374 observations) might appear limited compared to the quantity suggested by previous simulations^34^. However, we use a narrow age range (about one-fiftieth to one-third in size), task activity, and longitudinal data, which would all modify simulation estimates downwards. For example, growth charts using longitudinal data require much less data than those with cross-sectional data^15,17^. Together, these design choices increase the generalizability of our curves when using smaller samples. Though we left sites out when building our growth curves, our models were not fully externally validated. Lastly, while we demonstrate these growth charts’ utility through associations with prenatal exposures and postnatal outcomes, additional research is needed to assess widerspread utility, sensitivity in early life, and associations with longer-term neurocognitive outcomes. Integration of these growth charts into HAPPE facilitates such wider future testing.

## Conclusion

We demonstrate the feasibility and utility of task-derived brain function growth charts using the EEG-derived visual-evoked potential in awake infants from cohorts spanning three continents. The growth charts exhibit high levels of fit, stability within individual trajectories, and generalize well across different global cultural, linguistic, and socioeconomic contexts. These normative models are sensitive to environmental factors and emerging cognitive development and increase power to discover new results. Overall, we demonstrate the potential of the brain function growth charts in awake infants as an impactful tool to study neurodevelopment at scale.

## Methods

### Participants

All study procedures were approved by the relevant university Health Research Ethics Committees (Protocol Nos.: Khula - HREC REF 666/2021, Germina - CAAE 49671221.2.0000.0068, LABS - P00039731, GABA - P00028129). Informed consent was collected from caregivers on behalf of their infants. Cohort characteristics are summarized in Table 1, and demographic information by cohort in Table S1.

#### Cohort Descriptions

##### Khula Cohort from South Africa

Infants from the Khula South Africa cohort^49^ were recruited from the Gugulethu Midwife Obstetrics Unit, an antenatal clinic in an informal settlement, Gugulethu, in the Western Cape province of South Africa, as part of the Wellcome LEAP 1kD project. Data collection occurred at the University of Cape Town’s Neuroscience Institute. 97.46% of caregivers from this site spoke Xhosa as their primary language. 82.30% of caregivers included in this analysis were recruited prenatally, with the remaining 17.70% being recruited shortly after birth. Women were eligible to be enrolled in the study if they were (i) pregnant, (ii) in their third trimester of pregnancy (28-36 weeks) or up to three months postpartum, and (iii) over the age of 18 years at the time of recruitment. Inclusion criteria for the study included (a) singleton pregnancy, (b) no psychotropic drug endorsement during pregnancy, (c) no infant congenital malformation or abnormalities (e.g., spina bifida, Down’s syndrome), and (d) no significant delivery complications (e.g., uterine rupture, birth asphyxia). Infants from this cohort contributed Visual-Evoked Potential EEG, cognitive outcome (i.e., GSED-LF), and prenatal alcohol exposure data to this paper.

##### Germina Cohort from Brazil

Infants from the Germina cohort^48^ were recruited from São Paulo, Brazil, as part of the Wellcome LEAP 1kD project. Data collection occurred at the University of São Paulo Medical School. 100% of caregivers from this site spoke Brazilian Portuguese, and assessments were administered in Brazilian Portuguese. Infants included in this analysis were recruited postnatally, with the average age at recruitment being 2.4 months. Inclusion criteria were (a) maternal age of 20-45 years, (b) infant age of 3 months 0 days to 3 months and 29 days, (c) gestational age of >37 weeks, (d) infant birth weight >2000 grams, (e) no substance endorsement during pregnancy, (f) no history of severe maternal mental disorders (e.g., psychosis, bipolar disorder), (g) no delivery complications requiring medical intervention (e.g., perinatal asphyxia, shoulder dystocia, excessive bleeding), (h) no infant genetic syndrome or auditory/visual impairment diagnoses, and (i) availability for in-person lab visits. Infants from this cohort contributed visual-evoked potential EEG data to this paper.

##### LABS Cohort from USA

Infants from the LABS cohort were recruited from the greater Boston area in the United States of America as part of the Wellcome LEAP 1kD project. Data collection occurred at Boston Children’s Hospital. Based on inclusion criteria, English was the primary language for all caregivers from this site. Infants included in this analysis were recruited postnatally. The average age at recruitment varies by wave, with waves beginning at 3 months (+/- 7 days), 6 months (+/- 14 days), 12 months (+/- 14 days), 18 months (+/- 28 days), and 24 months (+/- 42 days). Inclusion criteria were (a) English was the primary language spoken at home (greater than 75% of the language spoken), (b) parental age was 18 years or older, (c) gestational age of 36-40 weeks, (d) infant birth weight of 2500-4500 grams, (e) no prenatal, postnatal, or birth-related complications, (f) no diagnosis of genetic, metabolic, syndromic, or neurological disorders, (g) no hearing or vision problems (unless corrected). Infants from this cohort contributed visual-evoked potential EEG data to this paper.

##### GABA Cohort from USA

Infants from the GABA cohort^47^ were recruited from the greater Boston area in the United States of America as part of a larger study investigating the effects of early general anesthesia on brain development, and data collection occurred at Boston Children’s Hospital. 92.06% of caregivers from this site reported speaking English with their child. All infants included in this analysis were recruited postnatally. Inclusion criteria included (a) parents/legal guardians ≥18 years old without developmental disabilities, without being wards of the state and willing to complete all study visits, (b) infant age ≤280 postnatal days, (c) gestational age of ≥32 weeks, (d) lack of confirmed or suspected clinical seizures, (e) lack of confirmed or suspected auditory or visual impairment, (f) no diagnosis of a genetic disorder affecting development, (g) no ongoing medical condition that would interfere with the conduct and assessment of the study, and (h) fluent English speakers, in which English is spoken to the infant ≥75% awake hours. Infants recruited for the comparison group without general anesthetic exposure or hospital admittance and who opted to have their data used in future studies were included in this analysis. Infants from this cohort contributed visual-evoked potential EEG data to this paper.

## Measures

### Electroencephalography (EEG)

#### EEG Acquisition

At all sites, EEG data were acquired from infants seated in their caregiver’s lap in a dimly lit, quiet room. A 128-channel high-density HydroCel Geodesic Sensor Net (EGI, Eugene, OR), amplified with a NetAmps 400 high-input amplifier (GABA cohort used the 300 series) and recorded via an Electrical Geodesics, Inc. (EGI, Eugene, OR) system was used. A 1000 Hz sampling rate was used at the South Africa and U.S.A. sites. A 500 Hz sampling rate was used at the Brazil site. EEG data were online referenced to the vertex (channel Cz) through the EGI Netstation software. Impedances were kept below 100KΩ following the impedance capabilities of the high-impedance amplifiers. For the Khula cohort, Geodesic Sensor Nets with modified tall pedestals designed to improve the inclusion of infants with thick/curly/tall hair were used as needed across participants. Shea moisture leave-in castor oil conditioner with insulating ingredients was applied to hair across the scalp before net placement to improve impedances and participant comfort^104^. At all sites, the Visual-Evoked Potential (VEP) task was presented using Eprime software (Psychology Software Tools, Pittsburgh, PA) on a desktop computer with an external monitor ranging from 19.5 to 21.5 inches on the diagonal facing the infant. The monitor was approximately 60-65 cm away from the infant. A standard phase-reversal VEP was induced with a black and white checkerboard (1cm x 1 cm squares within the board) stimulus that alternated presentation (black squares became white, white squares became black) every 500 milliseconds for a total of 100 trials (up to 200 trials in GABA cohort). Participant looking was monitored by video and by an assistant throughout data collection. The task was rerun if the participant looked away during the VEP task or was overly fussy. For the GABA cohort, stimulus presentation proceeded if a Tobii Eye Tracker system registered the infant’s gaze on the monitor or the experimenter could see the infant looking at the screen. For the Germina cohort, video recording of the experiment was manually reviewed. Periods when the infant looked away or was overly fussy were removed from the data.

#### EEG Data Pre-Processing

EEG data were exported from native Netstation .mff format to .raw format and then pre-processed using HAPPE v3.3 software (HAPPE v3.2 used for pre-processing in GABA, but updates between versions do not affect ERP measures), an automated open-source EEG processing software validated for infant data^80^. A subset of the 128 channels was selected for pre-processing, which excluded the rim electrodes as these are typically artifact-laden (channels excluded from pre-processing are included in Table S2). The HAPPE Pre-Processing pipeline was run with user-selected specifications outlined in Table S3. Parameters were largely consistent across sites except for line noise frequency, which differed by geographic region.

Pre-processed EEG data were considered usable and moved forward to VEP extraction if HAPPE pre-processing ran successfully, at least 20 trials were retained following bad trial rejection, and at least two good channels were kept within the visual ROI.

#### VEP Extraction

VEP waveforms were extracted and quantified using the HAPPE v3.3 Generate ERPs script from the HAPPE+ER pipeline^81^. Electrodes in the occipital region were selected as a region of interest (i.e., E70, E71, E75, E76, E83). The VEP waveform has three main components to be quantified: a negative N1 peak, a positive P1 peak, and a negative N2 peak. The features of interest for these components were peak latency and amplitude. Developmentally appropriate windows were selected for each age group with an overall N1 window range of 40 to 120 milliseconds, an overall P1 window range of 75 to 175 milliseconds, and an overall N2 window range of 100 to 325 milliseconds. Parameters (which were consistent across sites) used in extracting the ERPs with HAPPE+ER are summarized in Table S3.

All VEPs were visually inspected to ensure the automatically extracted values were correct and adjusted if observable peaks occurred outside the automated window bounds. Data quality for included files at each site is summarized in Table S4.

### Cognitive Outcomes

#### Global Scales for Early Development, Long Form (GSED-LF)

To investigate how deviation from the GAMLSS curves relates to emerging behavior, we collected the Global Scales for Early Development, Long Form (GSED-LF) from infants in the Khula cohort around age 428 days (*SD*=36.61). The GSED Long Form (GSED-LF) is a directly administered tool developed by the World Health Organization to collect population-level behavioral development at scale globally across diverse cultural and linguistic contexts^70–72^. The GSED-LF provides an overall “d-score” that represents overall cognitive, communication, and motor functioning holistically. The standardized daz-score was used as our cognitive outcome.

### Prenatal Alcohol Exposure

As previously described^51^, mothers in the Khula South Africa cohort self-reported alcohol endorsement during pregnancy using the Alcohol Exposure Questionnaire (AEQ)^105^ at the first research visit. We extracted the number of standardized alcohol units consumed per week in each trimester from the AEQ (number of drinking episodes per week in each trimester multiplied by number of drinks consumed per episode). Mothers also self-reported non-alcohol substance endorsement (i.e., tobacco, cannabis, cocaine, amphetamines, inhalants, sedatives/sleeping pills, hallucinogens, opioids, or other) in the three months before enrollment using the World Health Organization Alcohol, Smoking, and Substance Involvement Screening Test ^106^. Infants with prenatal alcohol *and* non-alcohol substance exposures were excluded from the analysis to isolate the effects of alcohol^51,107,108^. 46 infants provided usable data at 112 days (*SD* = 26.86 days) whose mothers reported use of alcohol but no other comorbid substances.

### Analyses

#### Generalized Additive Models for Location, Scale, and Shape

Growth curves were modeled using generalized additive models for location, scale, and shape (GAMLSS), a framework developed to model complex data distributions^50^. GAMLSS is an extension of generalized additive models that allows independent functions to fit up to four parameters of the response distribution. As a result, myriad distribution families can be captured by modeling the mean, standard deviation, skewness, and kurtosis. GAMLSS is the World Health Organization’s recommended method for developing growth curves due to its flexibility in modeling the relationship between age and higher-order moments^4^.

GAMLSS models were fit to model the relationship between age and individual VEP features. We used p-splines as the smoothing function for each parameter. Distributions were chosen for each feature by minimizing the total validation Bayesian Information Criterion (BIC) when training the model in two datasets and validating in the third held-out dataset. We compared the standard forms of the following 3-4 parameter distributions: generalized *t* (GT), power exponential (PE), power exponential 2 (PE2), Sinh-Arcsinh (SHASH), *t*-family (TF and TF2), Box-Cox Cole and Green (BCCGo), Box-Cox power exponential (BCPEo), Box-Cox *t* (BCTo), gamma family (GAF), generalized beta type 2 (GB2), generalized gamma (GG), Johnson’s S_U_ (JSUo), generalized inverse gamma (GIG), skew normal types 1 and 2 (SN1 and SN2), skew exponential power type 1 (SEP1), and skew *t* (ST1). Degrees of freedom were optimized using maximum likelihood during the distribution comparison. With the distributions chosen for each feature, we selected the optimal generalized Akaike criterion penalty *k* using the same process. All models were fitted using up to 30 iterations of the Rigby-Stasinopoulos (RS)^109^ algorithim and up to 20 iterations of the Cole-Green (CG)^110^ algorithm. To evaluate biological sex as a covariate, we refit models with all features with sex included as an additional smoothing term for the *μ* term (i.e., the location or mean parameter) and compared the total validation BIC across all dataset combinations.

Model fit was quantified within-dataset by assessing the correspondence of the centiles predicted by the model and the actual quantiles of each data point. Specifically, we first predict the z-scores from the training data values and then convert them to their corresponding centile value. Using these centile scores, we predict the feature value given each age value from the training set and compare these to the actual values from the data. We report the Pearson correlation and root mean squared error (RMSE). Models were also assessed for fit through worm plots and Q-statistics. To understand the change in each parameter as a function of age, we created term plots representing *mu* (location), *sigma* (scale), *nu* (shape, skewness, or degrees of freedom, depending on the distribution), and *tau* (kurtosis) of each model (if applicable) over age (Fig. S14-19). Additional model details are reported in Table S8. We used numerical derivation to identify peaks and troughs by predicting each term over the range of ages and by predicting the 50th percentile for each feature.

### Centile Stability

To assess the stability of centiles within individuals, we binned centile scores into “lines” as is typically done to measure movement on growth curves. This is done because movement at the extreme centiles is not equivalent to movement near the center. We binned centiles at 5%, 10%, 25%, 50%, 75%, 90%, and 95%. Stability was measured in several ways based on how many centile lines an individual crossed. The primary metric was distance per timepoint, calculated by taking the absolute value of centile change between timepoints and dividing it by the number of timepoints. We also measured the variance, standard deviation, and range.

#### Associations with Cognitive Outcomes

We associated deviation from the GAMLSS curves associated with the GSED-LF z-scores measured at 14 months in the South Africa site. The z-scores from each feature were summed to generate composite deviation scores for each subject. This analysis was restricted to the South Africa site, as only these participants had available the GSED-LF outcome measures. Participants were only included if they had viable data for all three sessions (*n*=90). Associations between deviation scores and cognitive outcomes were quantified with Pearson’s *r* correlation.

We used partial correlations to assess the impact of potential covariates, including sex, age at VEP measurement, and percent frames removed during EEG preprocessing. We also fit a linear regression model predicting cognitive outcomes from VEP features to determine the relative influence of each feature. We used a 1000-fold bootstrap to determine the included features and coefficients of the optimal model with confidence intervals. While the R^2^ of the resultant model was high, the results were unstable due to the limited sample size (Fig. S34).

#### Associations with Prenatal Alcohol Exposure

Finally, we show that deviation scores can replicate results from conventional analyses. We associated increased prenatal alcohol exposure during the first trimester with longer P1 latencies in the first 6 months of life. Because P1 is preceded by N1, P1 latency is commonly corrected by subtracting N1 latency. We followed this approach with the z-scores to create corrected P1 latency z-scores. Associations between these scores and alcohol consumption were assessed by Pearson’s *r* correlation.

## Data Availability

The EEG and behavioral data are not publicly available due to ethical concerns. Access to the data is subject to a data sharing agreement as described in the original data papers ^47–49^. The GAMLSS curves are publicly available through the HAPPE software and can be found at https://github.com/PINE-Lab/HAPPE.

## Code Availability

HAPPE can be found at https://github.com/PINE-Lab/HAPPE. The GAMLSS R package can be found at https://cran.r-project.org/web/packages/gamlss/index.html. Other associated code can be found at https://github.com/chrisclaycamp/VEP-GAMLSS.

## Acknowledgements

The authors thank the caregivers, families, and infants who made this work possible. The Khula South Africa Team tag includes the following authors: Lauren Davel, Bokang Methola, Donna Herr, Marlie Miles, Michal Zieff, Thandeka Mazubane and Zayaan Goolam Nabi. The Germina Team tag includes the following authors: Alline Cristina de Campos, André Fujita, Carla R. Taddei, Maria Rita Passos-Bueno, Patricia Beltrão-Braga. The LABS Team tag includes the following authors: Trevor Bissert and Berit Hartjen. The GABA Team tag includes the following authors: Siobhan Coffman, Isabelle Kim, Alice Tao, Ellen Underwood, and Maria Maloney. The 1kD EEG Working Group tag includes the following authors: Nicolò Pini, William Fifer, Sarah A. McCormick, Priyanka Ghosh, Cara Bosco, and Michal R. Zieff. The 1kD Machine Learning Working Group tag includes the following authors: Peter Wijeratne, Kevin S. Bonham, Daniel C. Alexander, and Guilherme Zainotti Miguel Fahur Bottino. This research is supported by the Wellcome LEAP 1kD Program (KAD, GVP, CAN, ES, DS, LJGD), Boston Children’s Hospital (Trailblazer Research Award to LC and Sara Page Mayo Endowment for Pediatric Pain Research, Treatment, and Education to CBB), and the World Premier International Research Center Initiative–International Research Center for Neurointelligence (TKH). The views expressed in this article are those of the authors only.

