## Supplemental Materials for "Growth charts of infant visual neurodevelopment generalize across global contexts"

**Table S1**

*Demographic Characteristics by Cohort (i.e., Khula South Africa [A], Germina [B], LABS [C], GABA [D]).*

**Table S1A**

*Khula South Africa Cohort Demographic Information.*

| | **Total (N=303)** |
|---|---|
| **Infant Biological Sex** | |
| Female | 151 (49.8%) |
| Male | 152 (50.2%) |
| **Primary Spoken Language** | |
| Xhosa | 295 (97.4%) |
| English | 3 (1.0%) |
| Sotho | 2 (0.7%) |
| Afrikaans | 1 (0.3%) |
| Ndebele | 1 (0.3%) |
| Zulu | 1 (0.3%) |
| **Maternal Educational Attainment** | |
| Completed Grade 6 (Standard 4) to Grade 7 (Standard 5) | 8 (2.6%) |
| Completed Grade 8 (Standard 6) to Grade 11 (Standard 9) i.e., high school without matriculating | 135 (44.6%) |
| Completed Grade 12 (Standard 10) i.e., high school | 116 (38.3%) |
| Part of university/ college/ post-matric education | 29 (9.6%) |
| Completed university/ college/ post-matric education | 15 (5.0%) |
| **Annual Household Income (South African Rand/ZAR)[a]** | |
| Less than R11,999 | 51 (16.8%) |
| R12,000 – R59,999 | 142 (46.9%) |
| R60,000 – R119,999 | 69 (22.8%) |
| More than R120,000 | 14 (4.6%) |
| Unknown | 27 (8.9%) |
| **Maternal Place of Birth** | |
| South Africa | 298 (98.3%) |
| In the African Continent (not South Africa) | 5 (1.7%) |

[a]At the time of writing (11/17/24), 1 USD=5.79399 Brazilian Real=18.1675 ZAR

**Table S1B**

*Germina Cohort Demographic Information.*

| | Total (N=390) |
|---|---|
| **Infant Biological Sex** | |
| Female | 192 (49.2%) |
| Male | 198 (50.8%) |
| **Primary Spoken Language** | |
| Portuguese Language | 387 (99.2%) |
| English Language | 1 (0.3%) |
| Spanish Language | 1 (0.3%) |
| Missing | 1 (0.3%) |
| **Maternal Educational Attainment** | |
| Completed 8th Grade | 4 (1.0%) |
| Completed High School | 75 (19.2%) |
| Completed Undergraduate Degree | 271 (69.5%) |
| Completed Graduate Degree (Master's or Doctorate) | 40 (10.3%) |
| **Annual Household Income (Brazilian Real/BRL)[a]** | |
| Less than R$27599 | 40 (10.3%) |
| R$27600 - R$59999 | 48 (12.3%) |
| R$60000 - R$119999 | 106 (27.2%) |
| R$120000 - R$239999 | 89 (22.8%) |
| Greater than R$240000 | 65 (16.7%) |
| Missing | 42 (10.8%) |
| **Infant Ethnicity** | |
| White | 260 (66.7%) |
| Pardo | 93 (23.8%) |
| Black | 21 (5.4%) |
| East Asian Descent | 16 (4.1%) |

[a]At the time of writing (11/17/24), 1 USD=5.79399 Brazilian Real=18.1675 ZAR

**Table S1C**

*LABS Cohort Demographic Information.*

| | Total (N=56) |
|---|---|
| **Infant Biological Sex** | |
| Female | 27 (48.2%) |
| Male | 29 (51.8%) |
| **Primary Spoken Language** | |
| English | 48 (85.7%) |
| Spanish | 2 (3.6%) |
| Equal Distribution of English and Spanish | 2 (3.6%) |
| Other | 1 (1.8%) |
| Unknown | 3 (5.4%) |
| **Caregiver 1 Educational Attainment** | |
| High school/GED | 2 (3.6%) |
| Associate's Degree | 6 (10.7%) |
| Bachelor's Degree | 15 (26.8%) |
| Master's Degree | 20 (35.7%) |
| M.D., Ph.D., J.D. or equivalent | 11 (19.6%) |
| Unknown/Do not wish to Disclose/Missing | 2 (3.6%) |
| **Annual Household Income (United States Dollar/USD)**[a] | |
| Less than $49,999 | 3 (5.4%) |
| $50,000 - $99,999 | 10 (17.9%) |
| $100,000 - $149,999 | 6 (10.7%) |
| $150,000 - $199,999 | 11 (19.6%) |
| Greater than $200,000 | 19 (33.9%) |
| Unknown/Do not wish to Disclose/Missing | 7 (12.5%) |
| **Infant Ethnicity** | |
| Not Hispanic, Latino/a, or Spanish origin | 42 (75%) |
| Unspecified Hispanic, Latino/a, or Spanish origin | 4 (7.1%) |
| Mixed Hispanic, Latino/a, or Spanish origin | 3 (5.4%) |
| Mexican, Mexican American, Chicano/a | 2 (3.6%) |
| Puerto Rican | 2 (3.6%) |
| Unknown | 3 (5.4%) |
| **Infant Race** | |
| White | 33 (58.9%) |
| Multiracial | 14 (25%) |
| Black or African American | 2 (3.6%) |
| Asian American | 1 (1.8%) |
| East, South Asia, Southeast Asia (Asian) | 1 (1.8%) |
| Do not wish to Disclose/Missing | 5 (8.9%) |

[a]At the time of writing (11/17/24), 1 USD=5.79399 Brazilian Real=18.1675 ZAR

**Table S1D**

*GABA Cohort Demographic Information.*

| | Total (N=53) |
|---|---|
| **Infant Biological Sex** | |
| Female | 19 (35.8%) |
| Male | 34 (64.2%) |
| **Primary Spoken Language** | |
| English | 50 (94.3%) |
| Chinese | 1 (1.9%) |
| French | 1 (1.9%) |
| German | 1 (1.9%) |
| **Maternal Educational Attainment** | |
| High school/GED | 2 (3.8%) |
| Associate's Degree | 2 (3.8%) |
| Bachelor's Degree | 19 (35.8%) |
| Master's Degree | 19 (35.8%) |
| M.D., Ph.D., J.D. or equivalent | 10 (18.9%) |
| Missing | 1 (1.9%) |
| **Annual Household Income (United States Dollar/USD)**[a] | |
| Less than $49,999 | 4 (7.6%) |
| $50,000 - $99,999 | 9 (16.9%) |
| $100,000 - $149,999 | 10 (18.9%) |
| $150,000 - $199,999 | 9 (17.0%) |
| Greater than $200,000 | 18 (34.0%) |
| Unknown | 1 (1.9%) |
| Missing | 2 (3.8%) |
| **Infant Ethnicity** | |
| Not Hispanic, Latino/a, or Spanish origin | 49 (92.5%) |
| Hispanic, Latino/a, or Spanish origin | 3 (5.7%) |
| Missing | 1 (1.9%) |
| **Infant Race** | |
| White | 43 (81.1%) |
| Multiracial | 5 (9.4%) |
| Asian | 2 (3.8%) |
| Black or African American | 2 (3.8%) |
| Missing | 1 (1.9%) |

[a]At the time of writing (11/17/24), 1 USD=5.79399 Brazilian Real=18.1675 ZAR

**Table S2.**
*HAPPE Pre-Processing Script Parameters.*

| | |
|---|---|
| **HAPPE Version** | v3.3<br>(GABA cohort: v3.2, version change did not impact parameters) |
| **Density** | High (>30 channels) |
| **Resting State or Task** | Task |
| **ERP Analysis** | Yes |
| **Acquisition Layout** | 128 channel EGI HydroCel Geodesic Sensor Net |
| **Channels** | All except E1, E8, E14, E17, E21, E25, E32, E38, E43, E44, E48, E49, E56, E63, E68, E73, E81, E88, E94, E99, E107, E113, E114, E119, E120, E121, E125, E126, E127, E128 |
| **Line Noise** | |
| **Line Noise Frequency** | Khula cohort: 50 Hz<br>Germina, LABS, and GABA cohorts: 60 Hz |
| **Line Noise Reduction Method** | CleanLine - Default |
| **Resample** | Off |
| **Filter** | |
| **Filter - Lowpass Cutoff** | 30 Hz |
| **Filter - Highpass Cutoff** | 0.3 Hz (Germina cohort: 0.1 Hz) |
| **Filter Type** | EEGLAB's FIR |
| **Bad Channel Detection** | On |
| **Bad Channel Detection Method** | Default |
| **Wavelet Thresholding** | Default |
| **Wavelet Threshold Rule** | Hard |
| **MuscIL** | Off |
| **Segmentation** | On |
| **Starting Parameter for Stimulus** | - 0.1 seconds |
| **Ending Parameter for Stimulus** | 0.5 seconds (GABA cohort: 0.4 seconds) |
| **Task Offset** | Khula cohort: 11 milliseconds<br>LABS cohort: 9 milliseconds |

| | |
|---|---|
| | GABA cohort: 45 milliseconds<br>Germina cohort: 15 milliseconds |
| **Baseline Correction** | On |
| **Baseline Correction Start** | - 100 milliseconds |
| **Baseline Correction End** | 0 milliseconds |
| **Interpolation** | Off (Germina cohort: On) |
| **Segment Rejection** | On |
| **Segment Rejection Method** | Amplitude criteria only |
| **Minimum Segment Rejection Threshold** | - 200 |
| **Maximum Segment Rejection Threshold** | 200 |
| **Segment Rejection based on All Channels or ROI** | ROI |
| **ROI Channels** | E70, E71, E75, E76, E83 |
| **Re-Reference Method** | Average |

**Table S3**
*HAPPE+ER Generate ERPs Script Parameters.*

| | |
|---|---|
| **HAPPE Version** | 3.3 |
| **Average or Individual Trials** | Average |
| **Channels of Interest** | E70, E71, E75, E76, E83 |
| **Bad Channels Included/Excluded** | Included |
| **Calculating ERP Values** | On |
| **Windows** | Overall Ranges Across All Ages:<br>Min 40-120 milliseconds<br>Max 75-175 milliseconds<br>Min 100-325 milliseconds |

**Table S4**

*EEG Quality Control Metrics by Site.*

| | South Africa (N=622) | Brazil (N=500) | USA (N=252) |
|---|---|---|---|
| **VEP Trial Retention** | | | |
| **Number of Collected VEP Trials** | | | |
| Mean (SD) | 100 (0.731) | 73.1 (20.8) | 103 (11.6) |
| Median [Min, Max] | 100 [82.0, 100] | 77.0 [20.0, 100] | 100 [80.0, 200] |
| **Number of Retained VEP Trials** | | | |
| Mean (SD) | 95.6 (11.4) | 63.3 (22.6) | 101 (14.4) |
| Median [Min, Max] | 100 [21.0, 100] | 65.5 [20.0, 100] | 100 [39.0, 196] |
| **Region of Interest (ROI) Channel Retention** | | | |
| **Number of ROI Channels Retained** | | | |
| Mean (SD) | 4.26 (0.894) | 4.60 (0.522) | 4.37 (0.848) |

| | South Africa (N=622) | Brazil (N=500) | USA (N=252) |
|---|---|---|---|
| Median [Min, Max] | 4.50 [2.00, 5.00] | 5.00 [2.00, 5.00] | 5.00 [2.00, 5.00] |
| **Correlation of Data Pre-v s. Post- Wavelet Thresholding (Pearson's r)** | | | |
| **At 5 Hz** | | | |
| Mean (SD) | 0.464 (0.239) | 0.393 (0.209) | 0.309 (0.310) |
| Median [Min, Max] | 0.506 [0.00532, 0.947] | 0.391 [0.0147, 0.892] | 0.166 [0.000127, 0.954] |
| **At 8 Hz** | | | |
| Mean (SD) | 0.434 (0.235) | 0.355 (0.190) | 0.288 (0.316) |
| Median [Min, Max] | 0.450 [0.00833, 0.931] | 0.350 [0.0132, 0.805] | 0.117 [0.000845, 0.947] |
| **At 12 Hz** | | | |
| Mean (SD) | 0.377 (0.217) | 0.348 (0.194) | 0.260 (0.325) |
| Median [Min, Max] | 0.401 [0.00477, 0.857] | 0.329 [0.0157, 0.795] | 0.0619 [0.000593, 0.954] |

| | South Africa (N=622) | Brazil (N=500) | USA (N=252) |
|---|---|---|---|
| **At 20 Hz** | | | |
| Mean (SD) | 0.436 (0.228) | 0.446 (0.204) | 0.304 (0.335) |
| Median [Min, Max] | 0.476 [0.00680, 0.854] | 0.458 [0.0172, 0.847] | 0.115 [0.000130, 0.952] |

Note. EEG data were pre-processed and VEPs were extracted using HAPPE+ER v3.3 software (v3.2 used for pre-processing of GABA cohort data), an automated open-source EEG processing software validated for infant data (Monachino et al., 2022).

**Table S5**

*Model Fits Averaged Across All Pairs of Sites.*

| | Within-dataset | | Held-out dataset | |
|---|---|---|---|---|
| **VEP Feature** | **Pearson *r* [95% CI]** | **RMSE** | **Pearson *r* [95% CI]** | **RMSE** |
| N1 Amplitude | *1.0 [.99-1.0]* | *.29* | *.99 [.99-1.0]* | *1.46* |
| P1 Amplitude | *.88 [.86-.89]* | *2.44* | *.85 [.82-.87]* | *2.67* |
| N2 Amplitude | *.99 [.99-1.0]* | *.67* | *.97 [.96-.97]* | *2.54* |
| N1 Latency | *.95 [.94-.96]* | *2.83* | *.95 [.94-.96]* | *5.66* |
| P1 Latency | *.69 [.64-.73]* | *13.12* | *.74 [.69-.78]* | *19.15* |
| N2 Latency | *.68 [.64-.72]* | *36.58* | *.71 [.66-.77]* | *36.79* |

**Table S6**

*Fits of Models Trained on All Sites.*

| | Within-dataset | |
|---|---|---|
| **VEP Feature** | **Pearson *r* [95% CI]** | **RMSE** |
| N1 Amplitude | *1.0 [1.0-1.0]* | *.25* |
| P1 Amplitude | *.87 [.85-.88]* | *2.57* |
| N2 Amplitude | *.99 [.99-.99]* | *.64* |
| N1 Latency | *.97 [.96-.97]* | *2.34* |
| P1 Latency | *.68 [.65-.71]* | *13.07* |
| N2 Latency | *.66 [.63-.70]* | *37.36* |

**Table S7**

*Quantile Residuals of Models Trained on All Sites.*

| VEP Feature | Mean | Variance | Coef. of skewness | Coef. of kurtosis | Filliben correlation |
|---|---|---|---|---|---|
| N1 Amplitude | *-0.0006411042* | *1.000384* | *-0.01051739* | *3.065981* | *0.9994753* |
| P1 Amplitude | *0.0002978088* | *0.9932231* | *-0.020952* | *2.966315* | *0.9992585* |
| N2 Amplitude | *-.00005277866* | *1.000571* | *0.01371232* | *2.899419* | *0.9996197* |
| N1 Latency | *-0.007189375* | *1.000569* | *-0.05502868* | *2.926535* | *0.9990406* |
| P1 Latency | *-0.0005036508* | *0.9691467* | *-0.1029502* | *3.097965* | *0.9987868* |
| N2 Latency | *0.00144525* | *1.00155* | *0.1809422* | *3.123041* | *0.9981535* |

**Table S8**

*Model Fits Averaged Across Cross-Validation Folds.*

| VEP Feature | Distribution | Generalized Akaike Information Criterion penalty *k* | Parameter | Link function | Represents | Association with age | Effective degrees of freedom |
|---|---|---|---|---|---|---|---|
| N1 Amplitude | Sinh-Arcsinh (SHASH) | 27 | $\mu$ | identity | location | Linear increase | 2.00 |
| | | | $\sigma$ | log | scale | Linear decrease | 2.00 |
| | | | $\nu$ | log | skewness | Linear decrease | 2.00 |
| | | | $\tau$ | log | kurtosis | Linear increase | 2.00 |
| P1 Amplitude | Johnson's $S_U$ (JSUo) | 6 | $\mu$ | identity | location | Linear decrease | 2.00 |
| | | | $\sigma$ | log | scale | Linear decrease | 2.00 |
| | | | $\nu$ | identity | skewness | Nonlinear increase | 3.32 |
| | | | $\tau$ | log | kurtosis | Linear decrease | 2.00 |
| N2 Amplitude | Johnson's $S_U$ (JSUo) | 21 | $\mu$ | identity | location | Linear increase | 2.00 |
| | | | $\sigma$ | log | scale | Linear decrease | 2.00 |
| | | | $\nu$ | identity | skewness | Linear increase | 2.00 |

| | | | | | | | |
|---|---|---|---|---|---|---|---|
| | | | $\tau$ | log | kurtosis | Linear decrease | 2.00 |
| N1 Latency | *t*-Family (TF) | 17 | $\mu$ | identity | location | Upward parabola | 3.35 |
| | | | $\sigma$ | log | scale | Linear decrease | 2.33 |
| | | | $\nu$ | log | degrees of freedom | Linear decrease | 2.00 |
| P1 Latency | Johnson's $S_U$ (JSUo) | 30 | $\mu$ | identity | location | Linear decrease | 2.00 |
| | | | $\sigma$ | log | scale | Linear increase | 2.00 |
| | | | $\nu$ | identity | skewness | Downward parabola | 4.12 |
| | | | $\tau$ | log | kurtosis | Linear increase | 2.00 |
| N2 Latency | Generalized Inverse Gaussian (GIG) | 23 | $\mu$ | log | location | Nonlinear decrease | 5.58 |
| | | | $\sigma$ | log | scale | Linear decrease | 2.00 |
| | | | $\nu$ | identity | shape | Linear decrease | 2.00 |

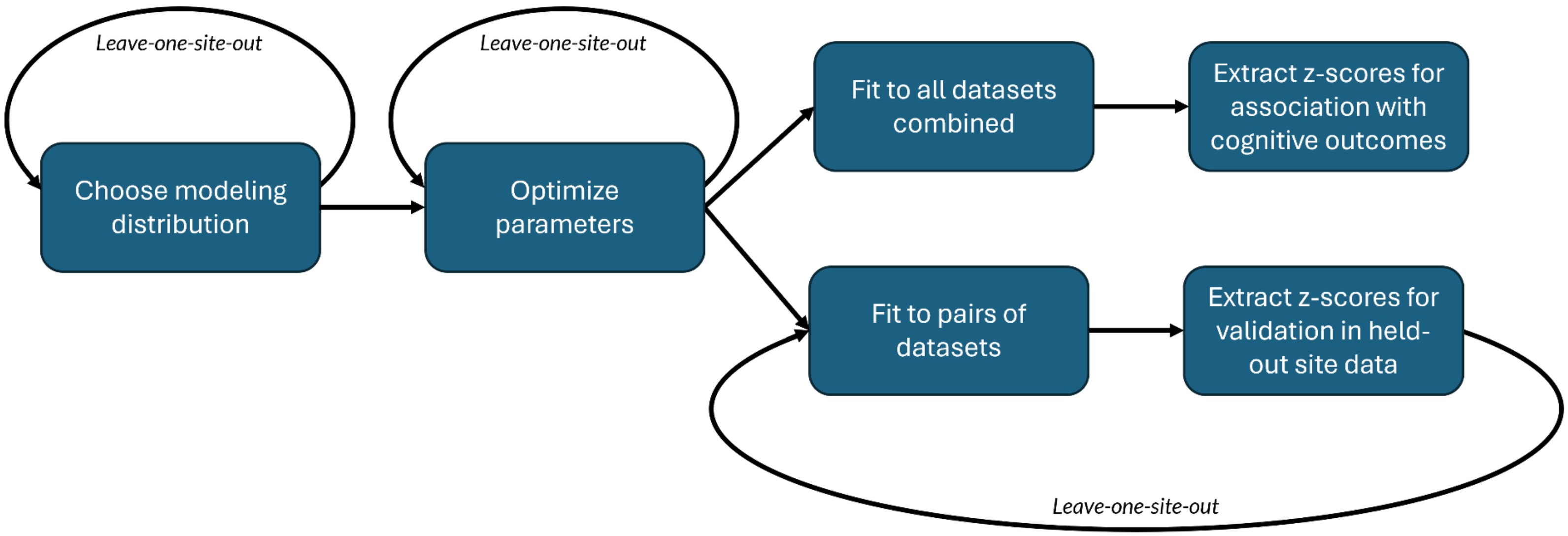

**Figure S1:** Methods flowchart for GAMLSS model fitting and tuning. For all *leave-one-site-out* loops, models were fit to each possible pair of site datasets and were validated in the held-out site dataset. Distributions and optimal smoothing parameters were chosen by minimizing the validation Bayesian Information Criterion over all loops.

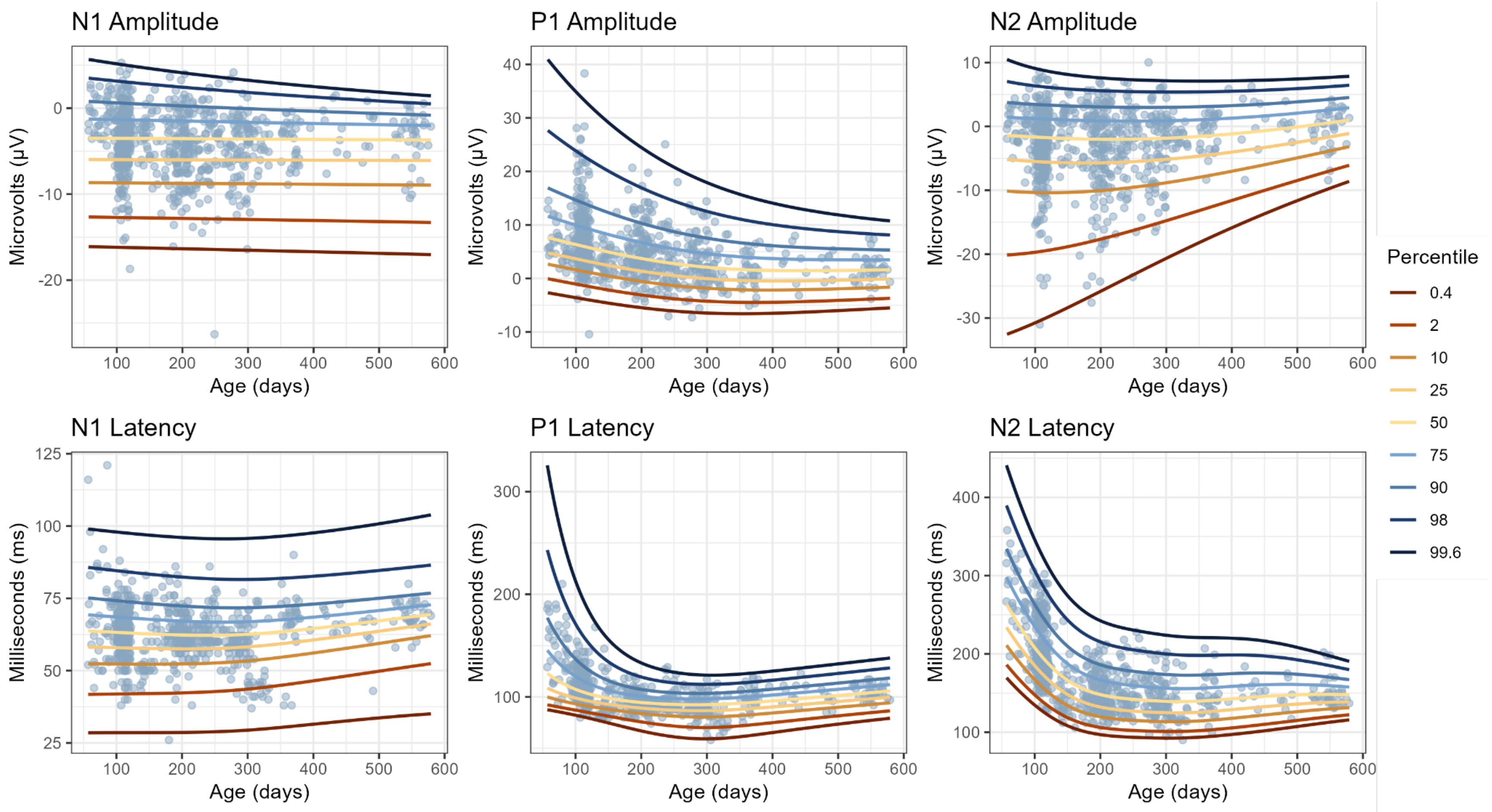

**Figure S2**: Centile growth curves trained on combined data from US and Brazil sites (*N* = 752 obs.) plotted on combined US and Brazil site data.

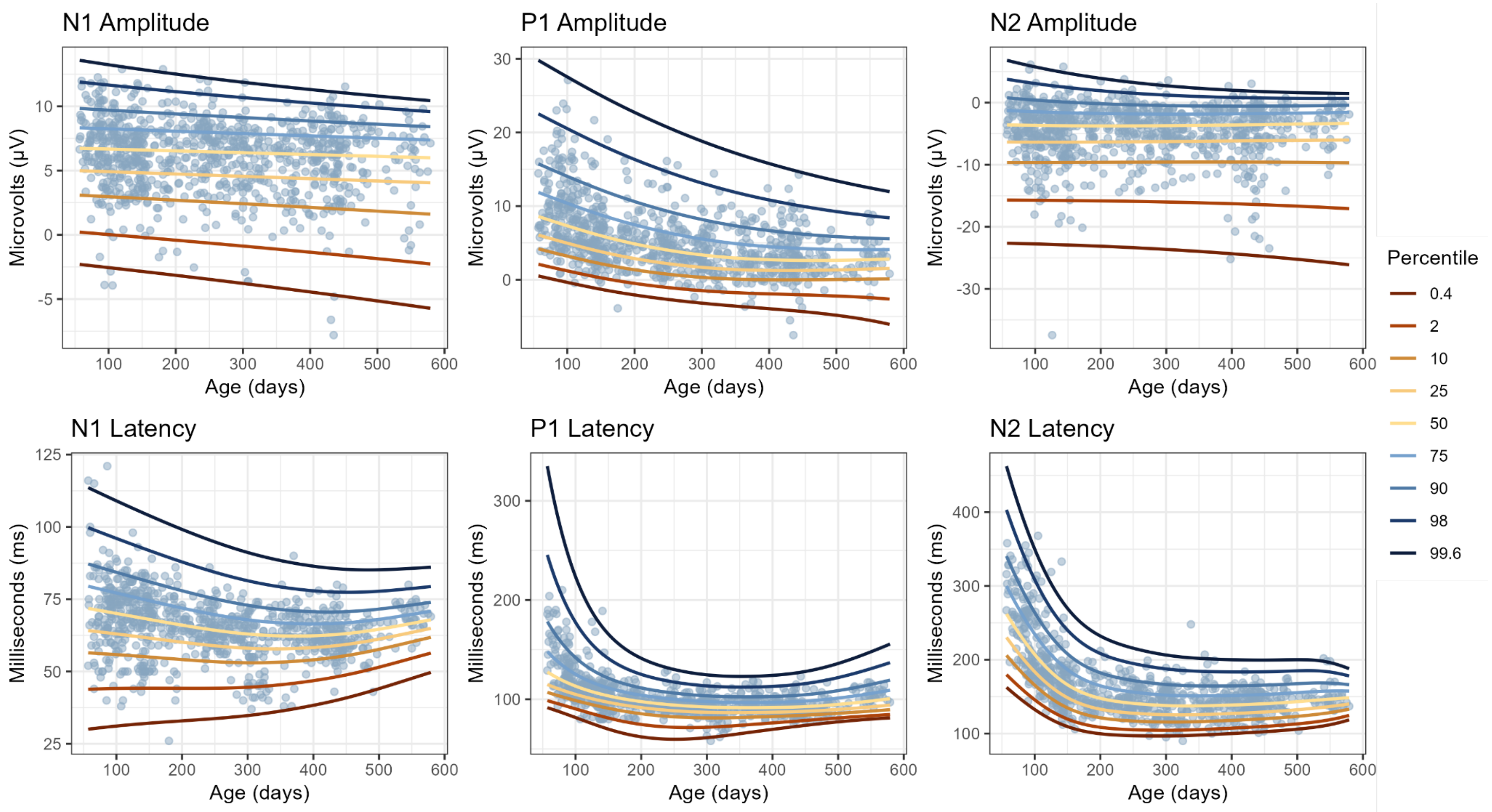

**Figure S3**: Centile growth curves trained on combined data from US and South Africa sites ($N$ = 871 obs.) plotted on combined US and South Africa site data.

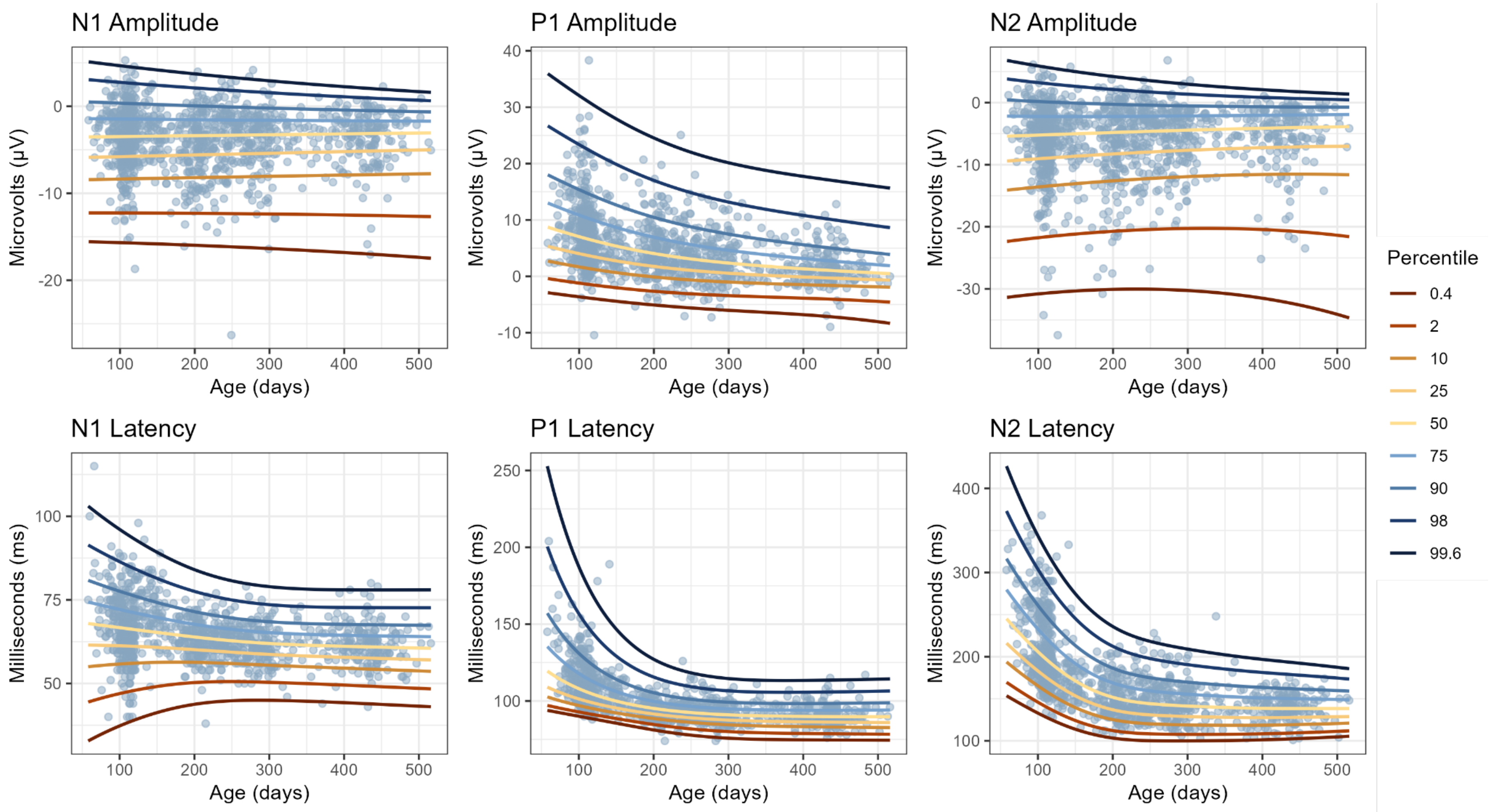

**Figure S4**: Centile growth curves trained on combined data from South Africa and Brazil sites (*N* = 1119 obs.) plotted on combined South Africa and Brazil site data.

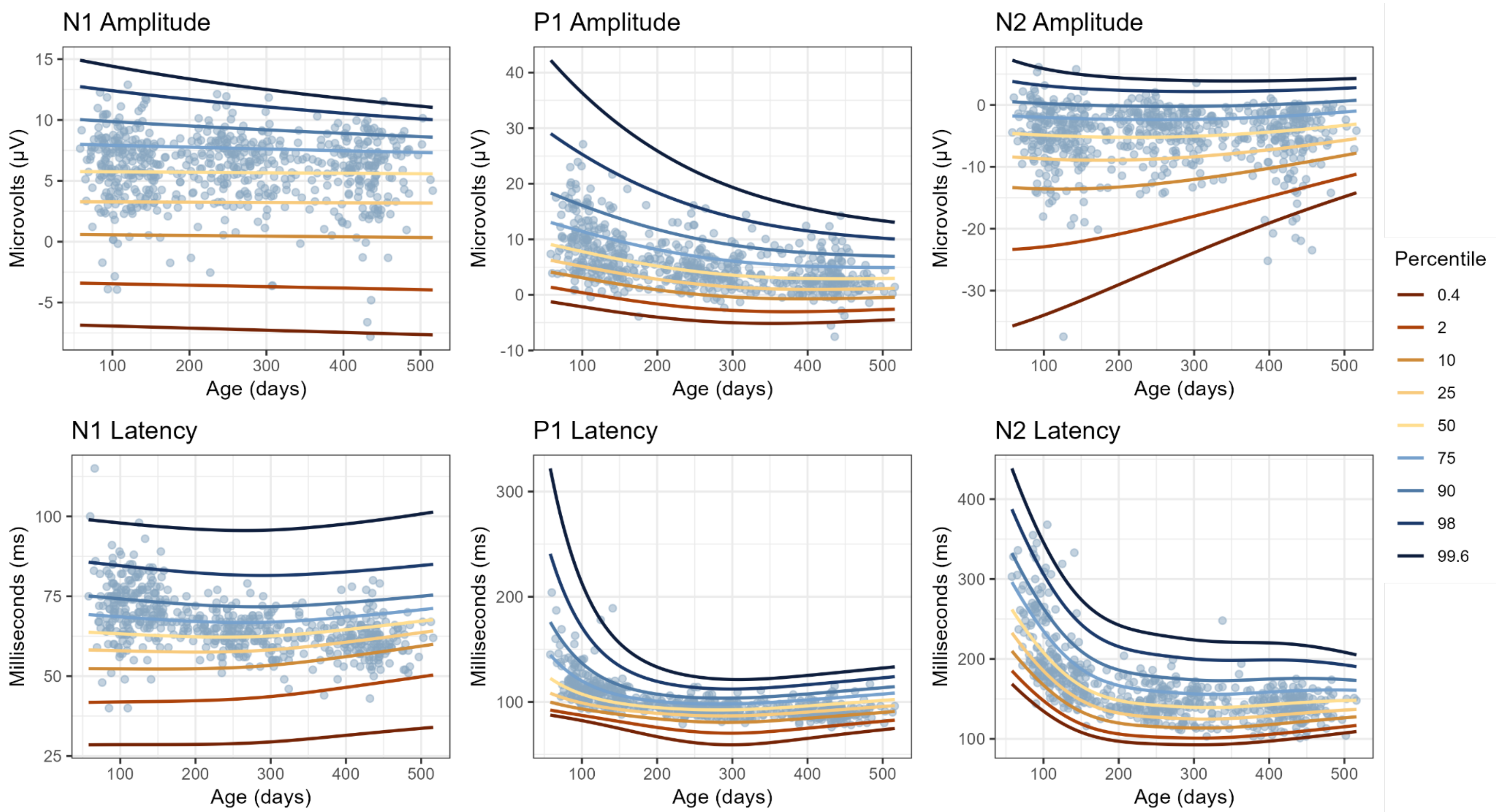

**Figure S5**: Centile growth curves trained on combined data from US and Brazil sites (*N* = 752 obs.) predicting held-out South Africa site data (*N* = 619 obs.).

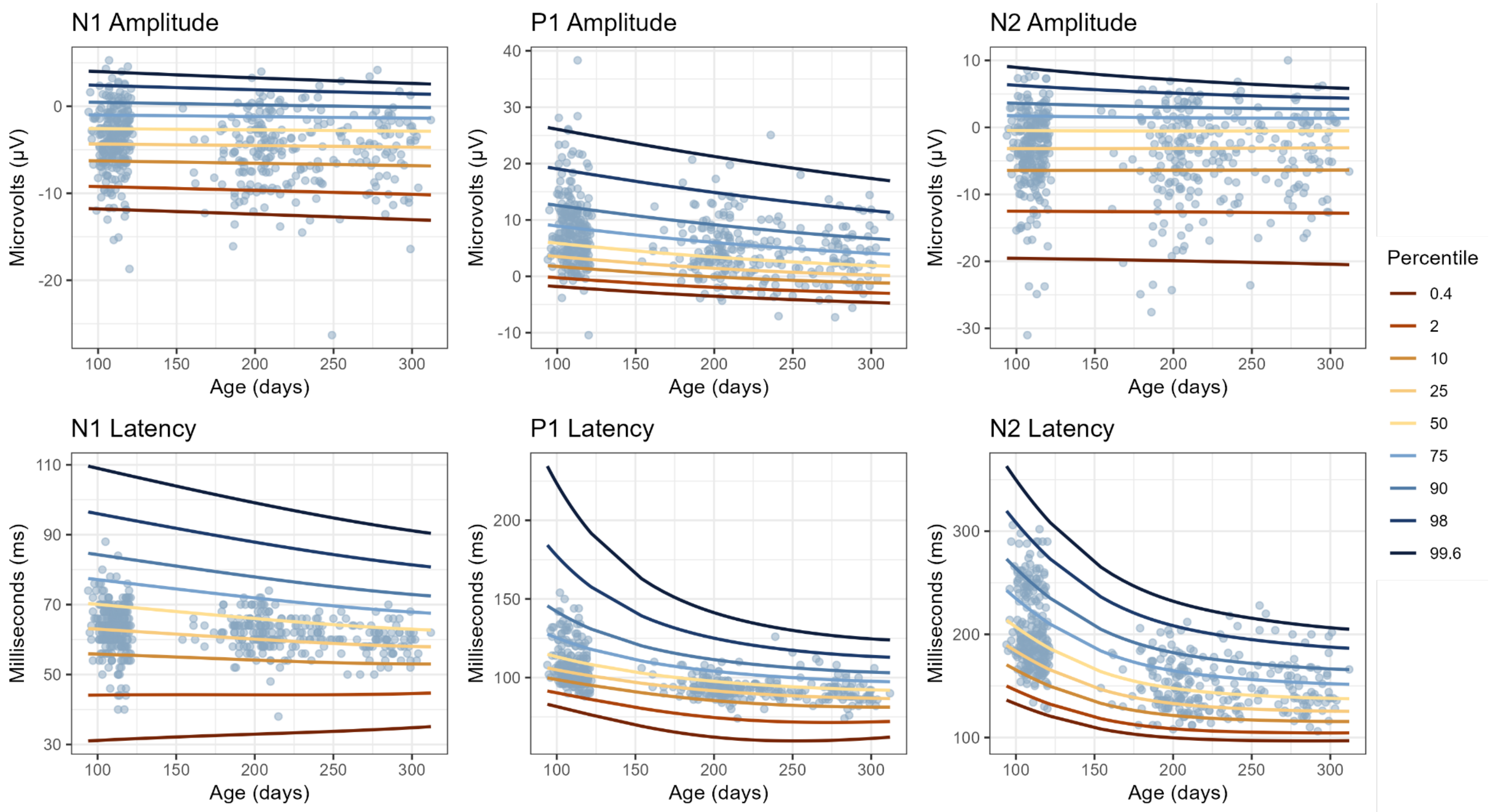

**Figure S6**: Centile growth curves trained on combined data from US and South Africa sites (*N* = 871 obs.) predicting held-out Brazil site data (*N* = 500 obs.).

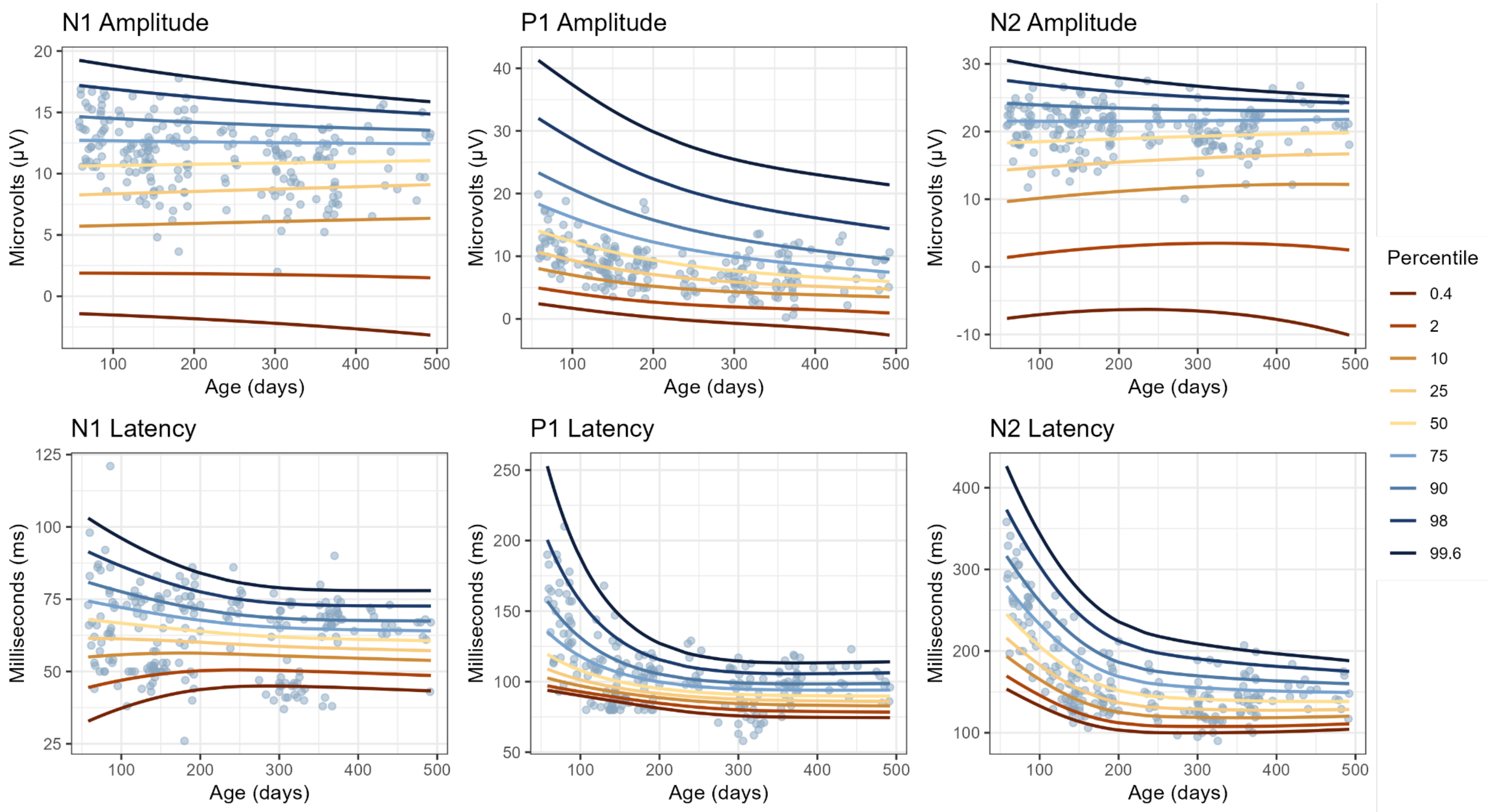

**Figure S7**: Centile growth curves trained on combined data from Brazil and South Africa sites ($N$ = 1119 obs.) predicting held-out US site data ($N$ = 215 obs.).

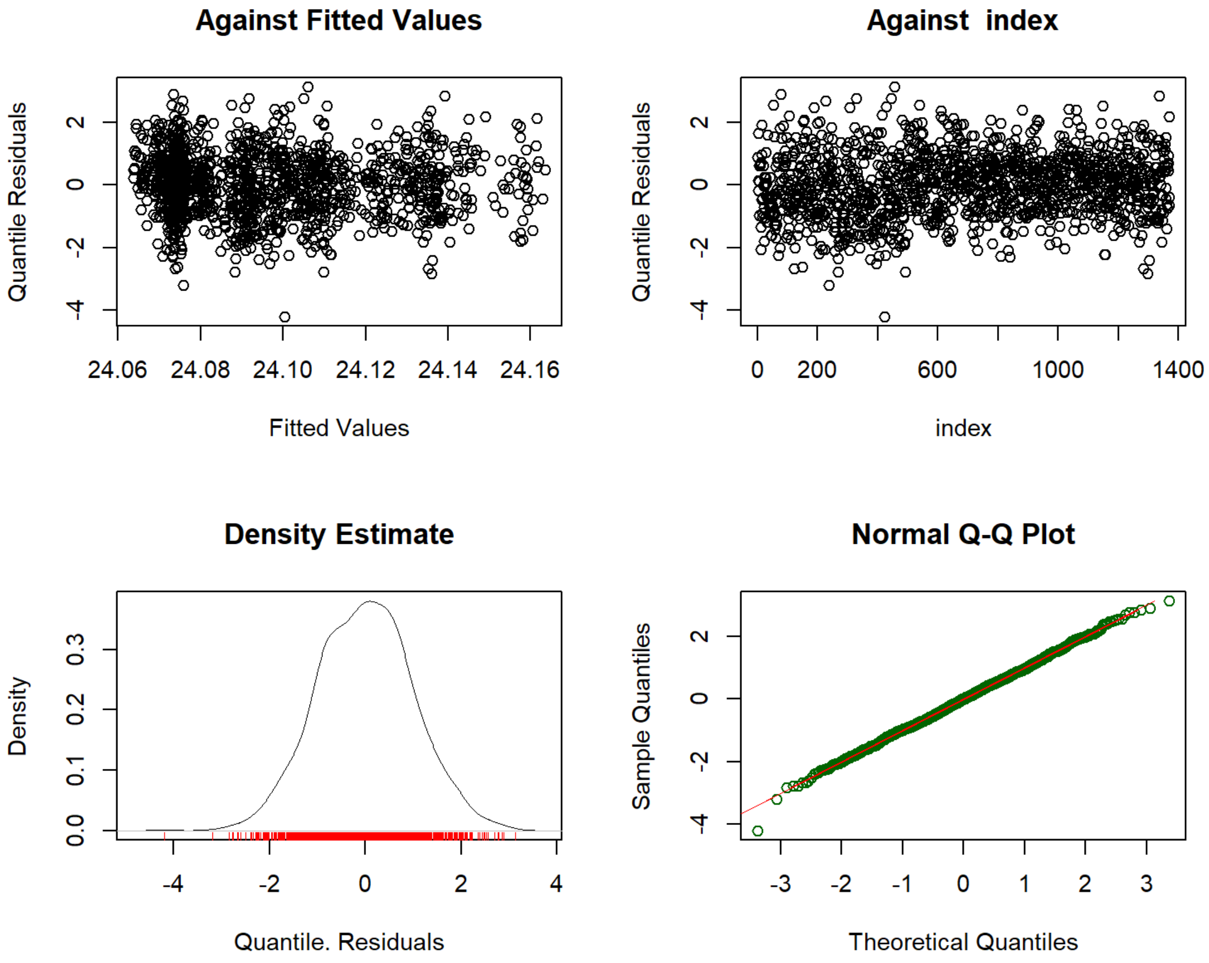

**Figure S8**: N1 Amplitude model residuals

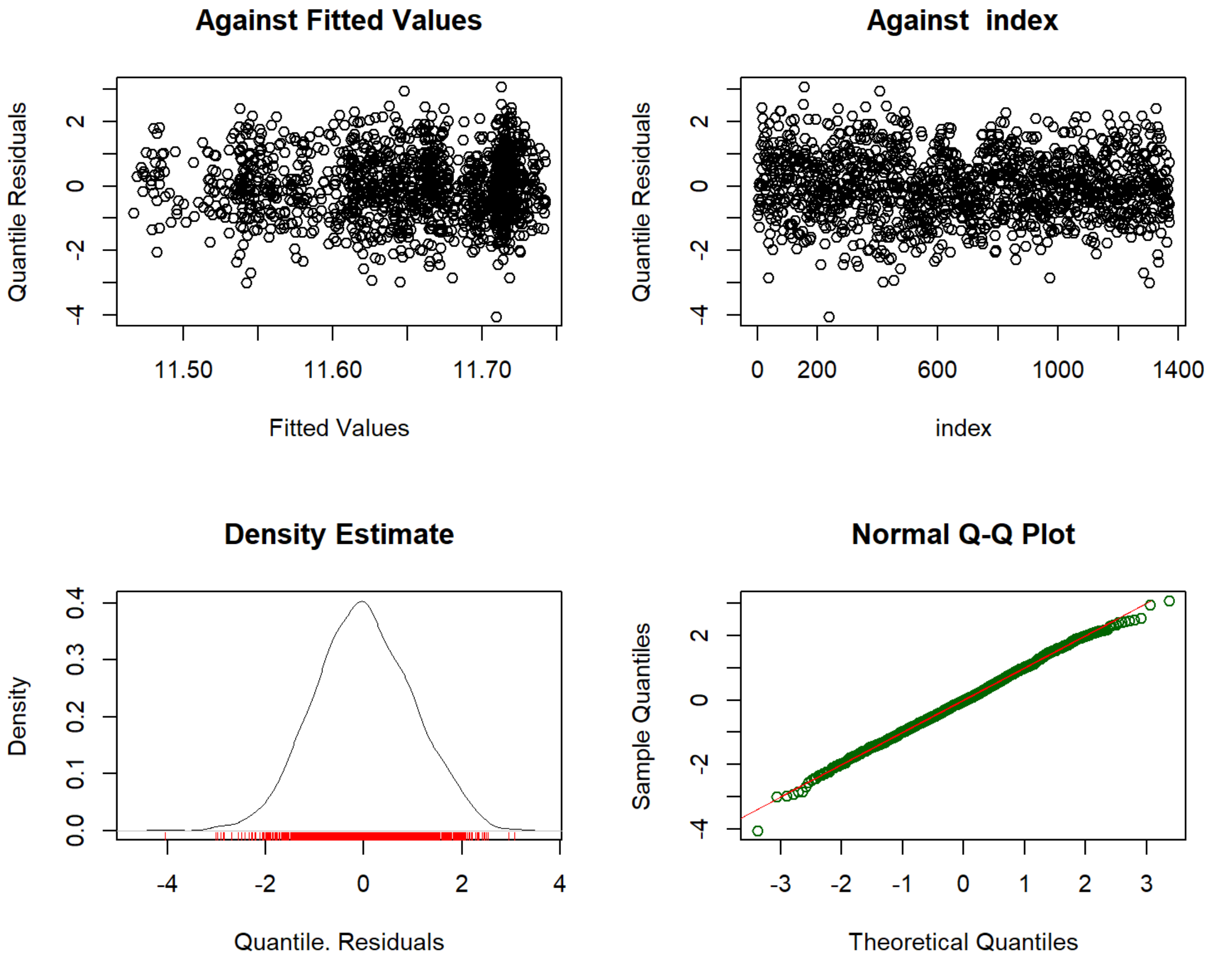

**Figure S9**: P1 Amplitude model residuals

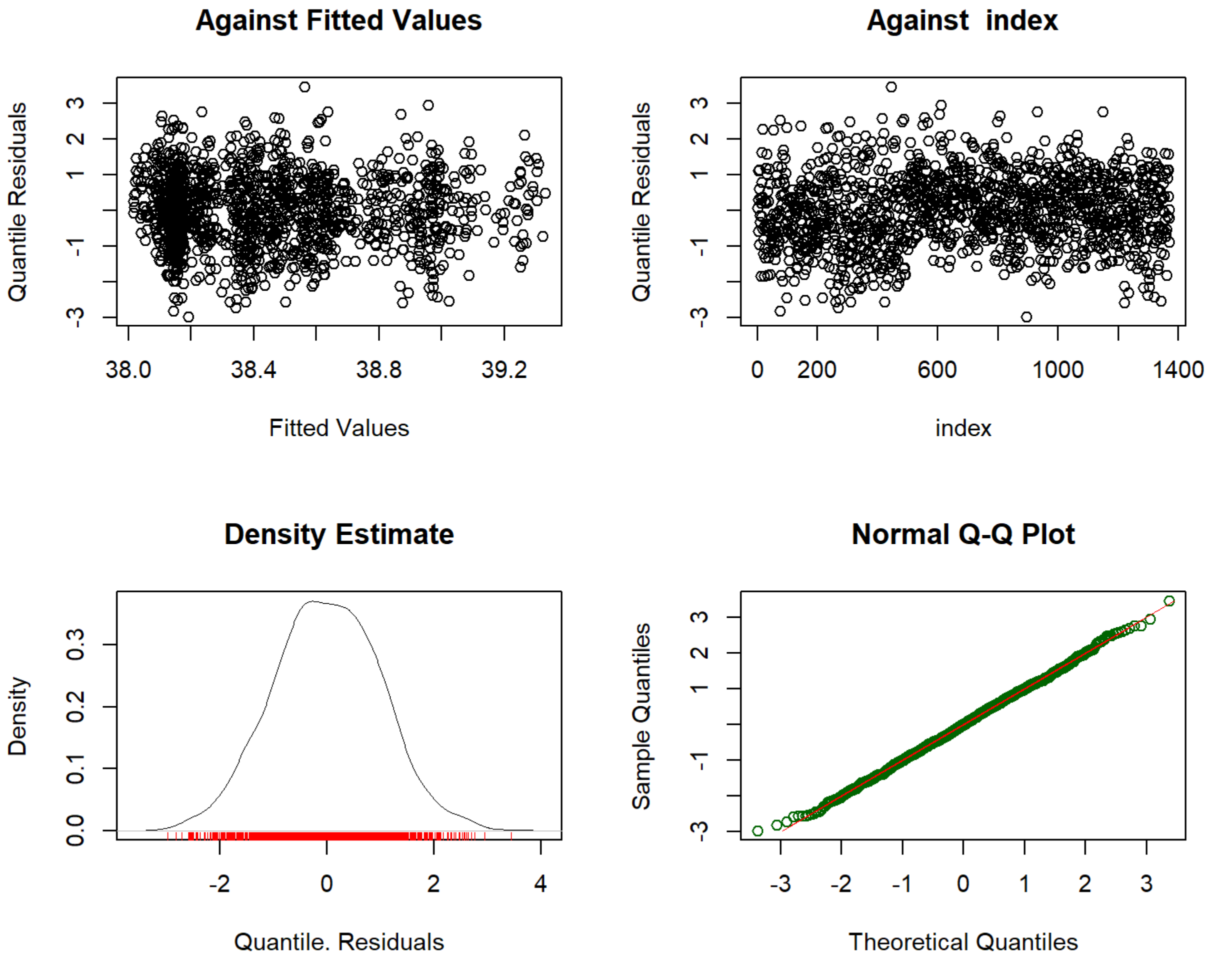

**Figure S10**: N2 Amplitude model residuals

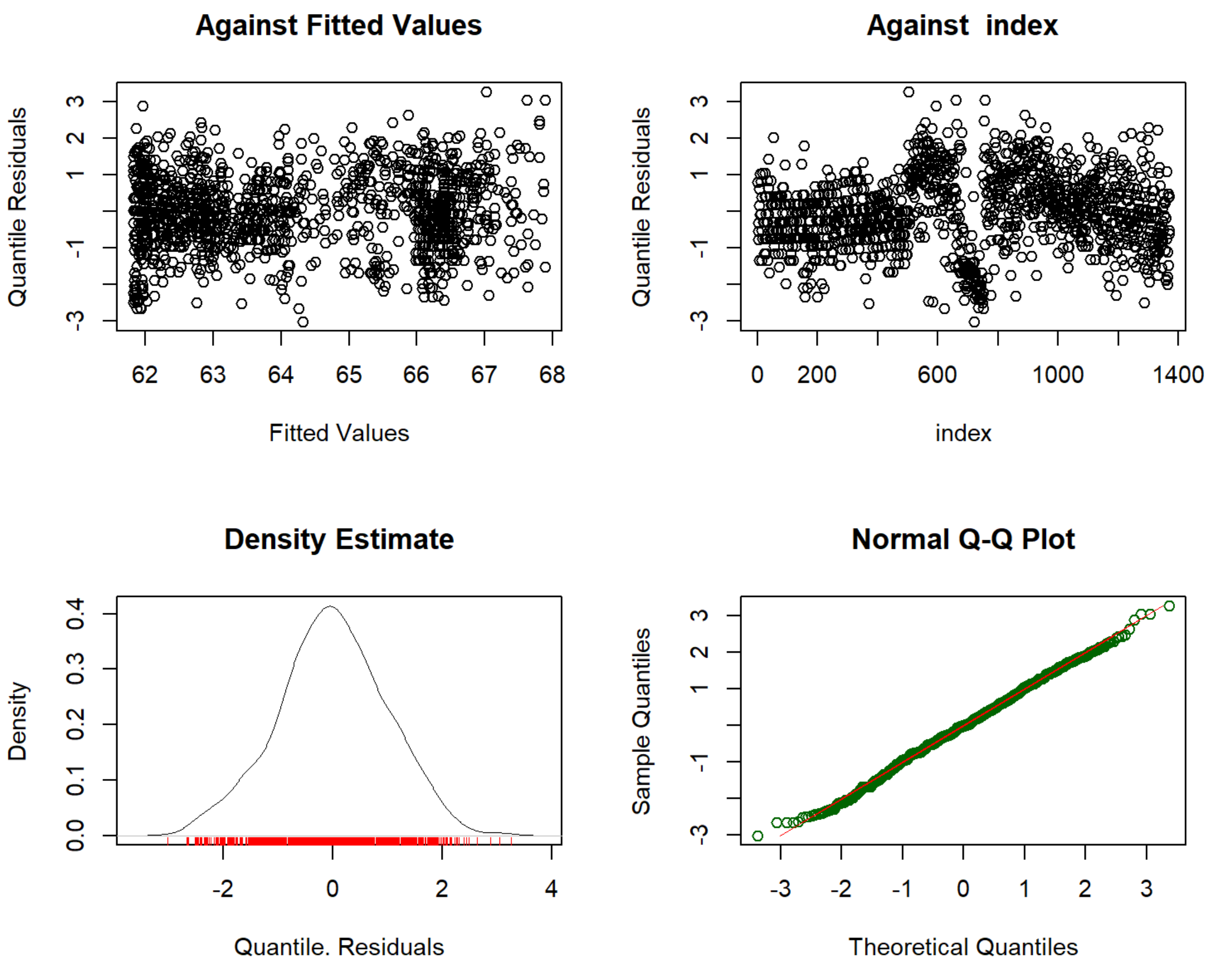

**Figure S11**: N1 Latency model residuals

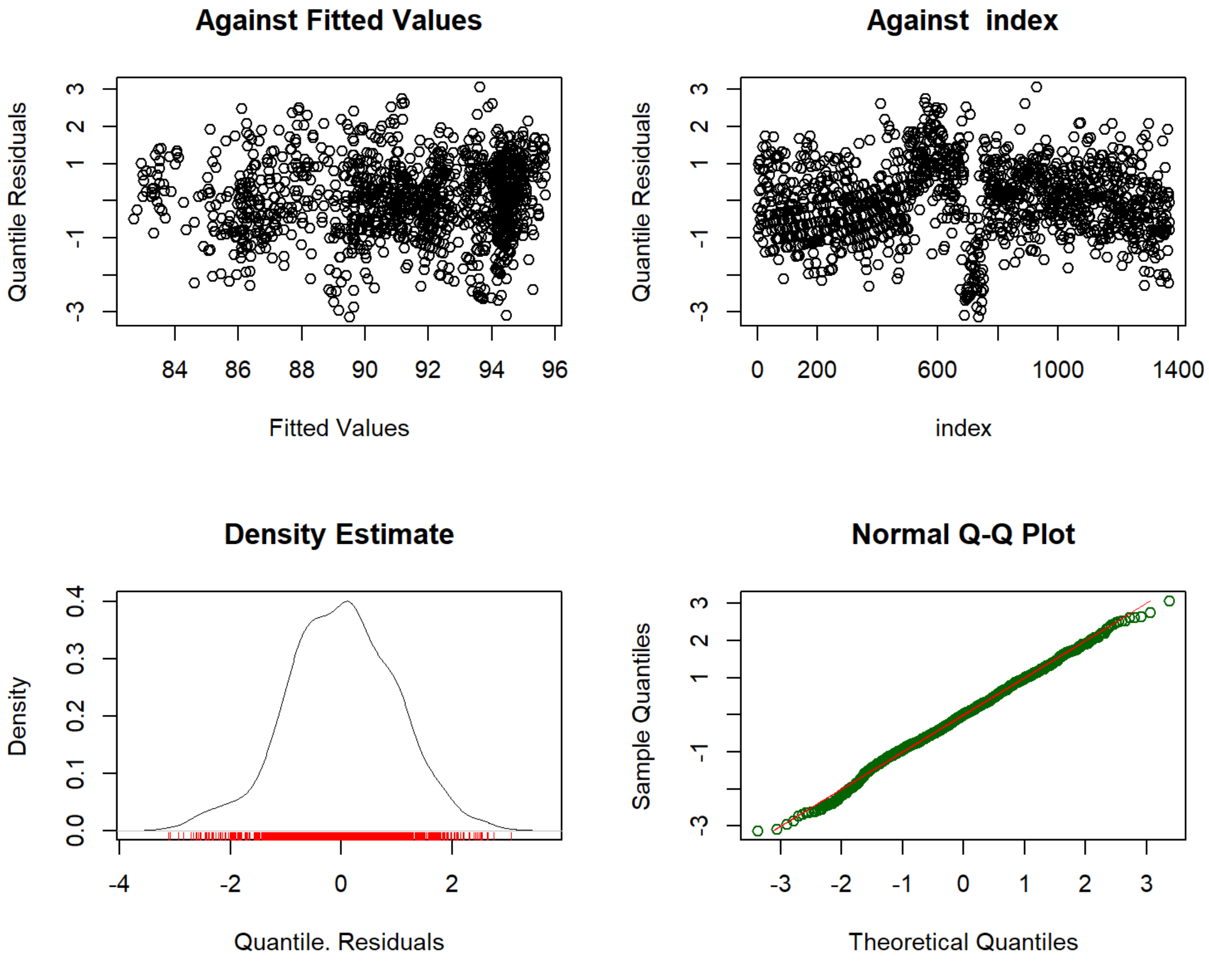

**Figure S12**: P1 Latency model residuals

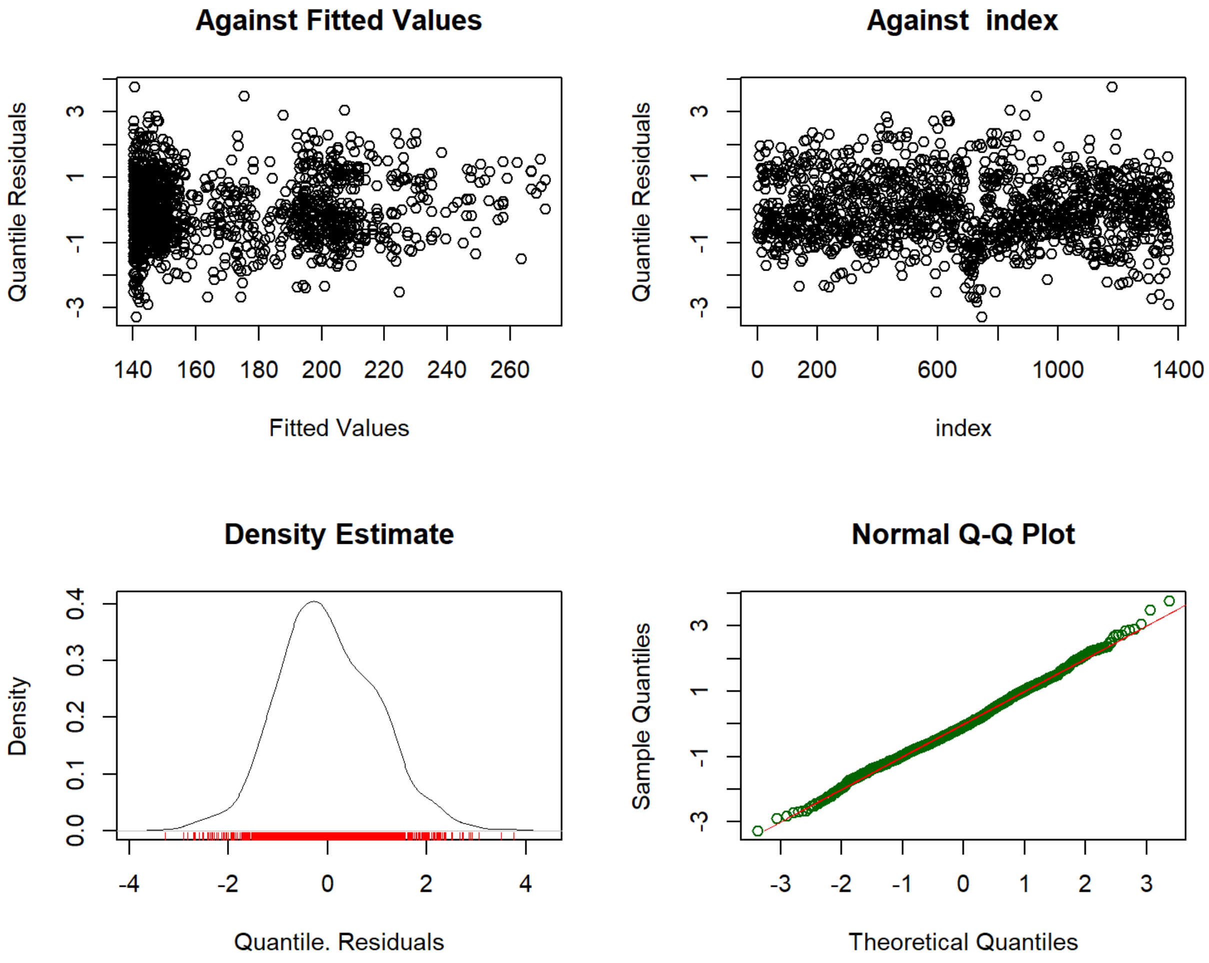

**Figure S13**: N2 Latency model residuals

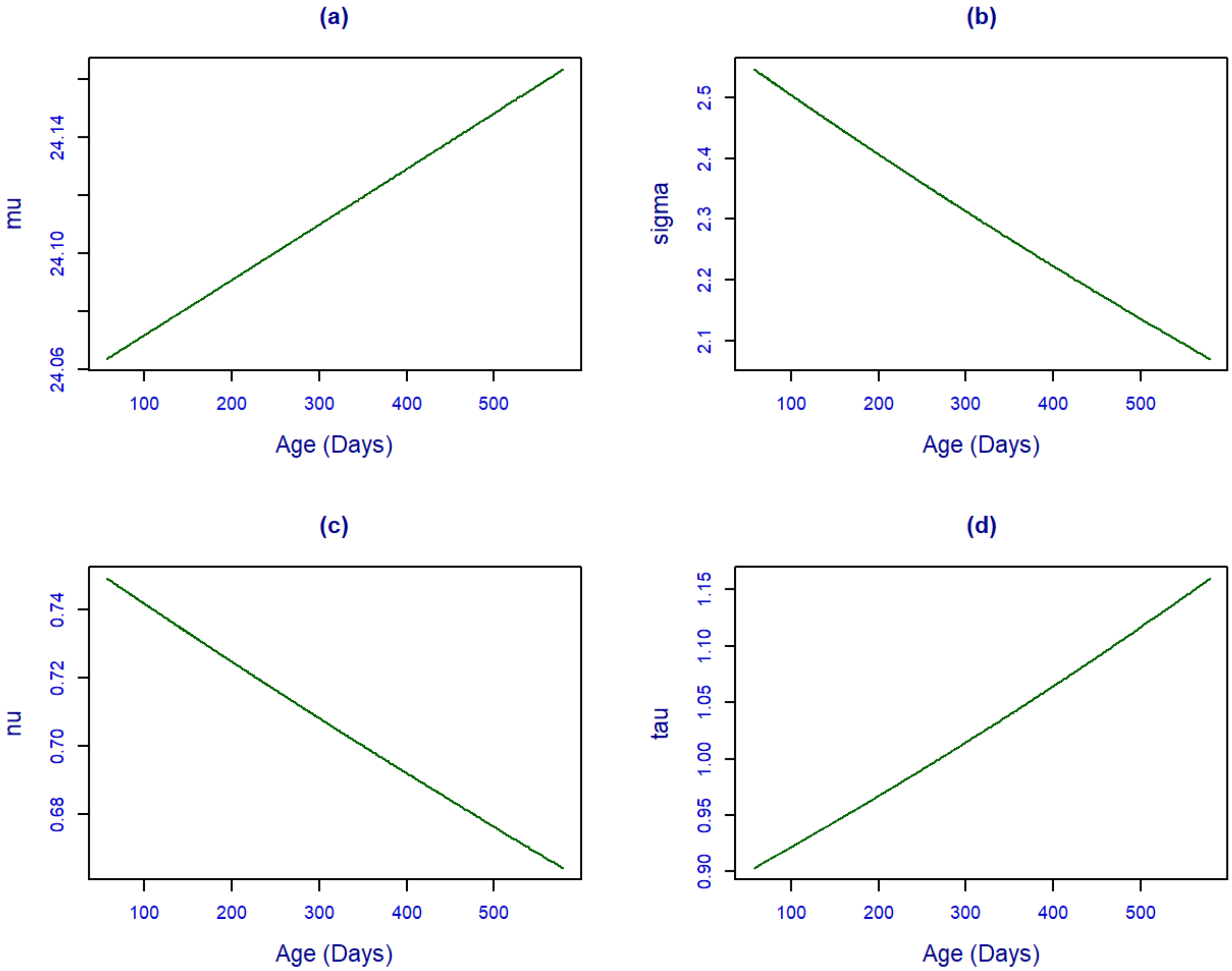

**Figure S14:** N1 Amplitude fitted term plots of relationship between age and a) mu, b) sigma, c) nu, and d) tau for the Sinh-Arcsinh growth curve model trained on all datasets.

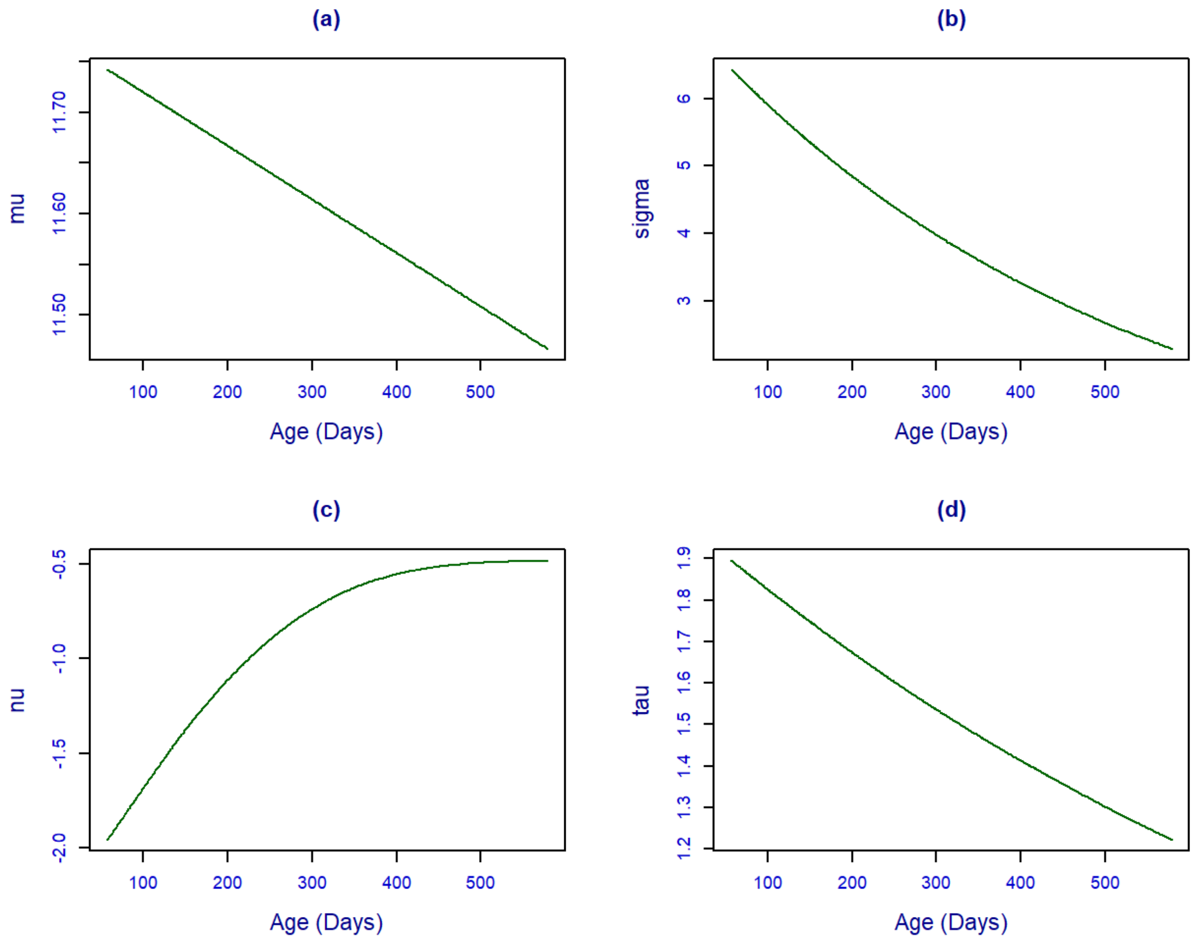

**Figure S15:** P1 Amplitude fitted term plots of relationship between age and a) mu, b) sigma, c) nu, and d) tau for the Johnson's $S_U$ growth curve model trained on all datasets.

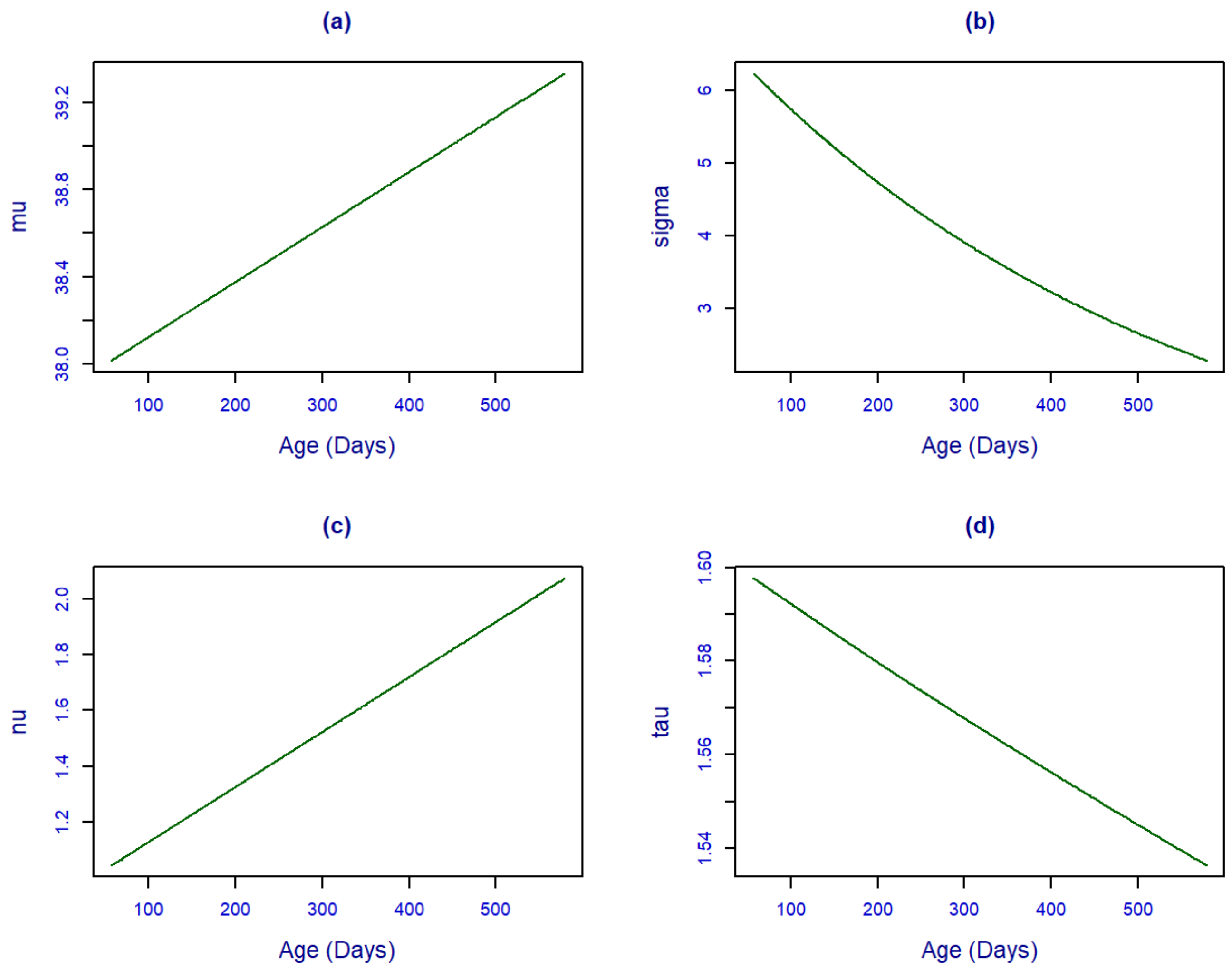

**Figure S16:** N2 Amplitude fitted term plots of relationship between age and a) mu, b) sigma, c) nu, and d) tau for the Johnson's $S_U$ growth curve model trained on all datasets.

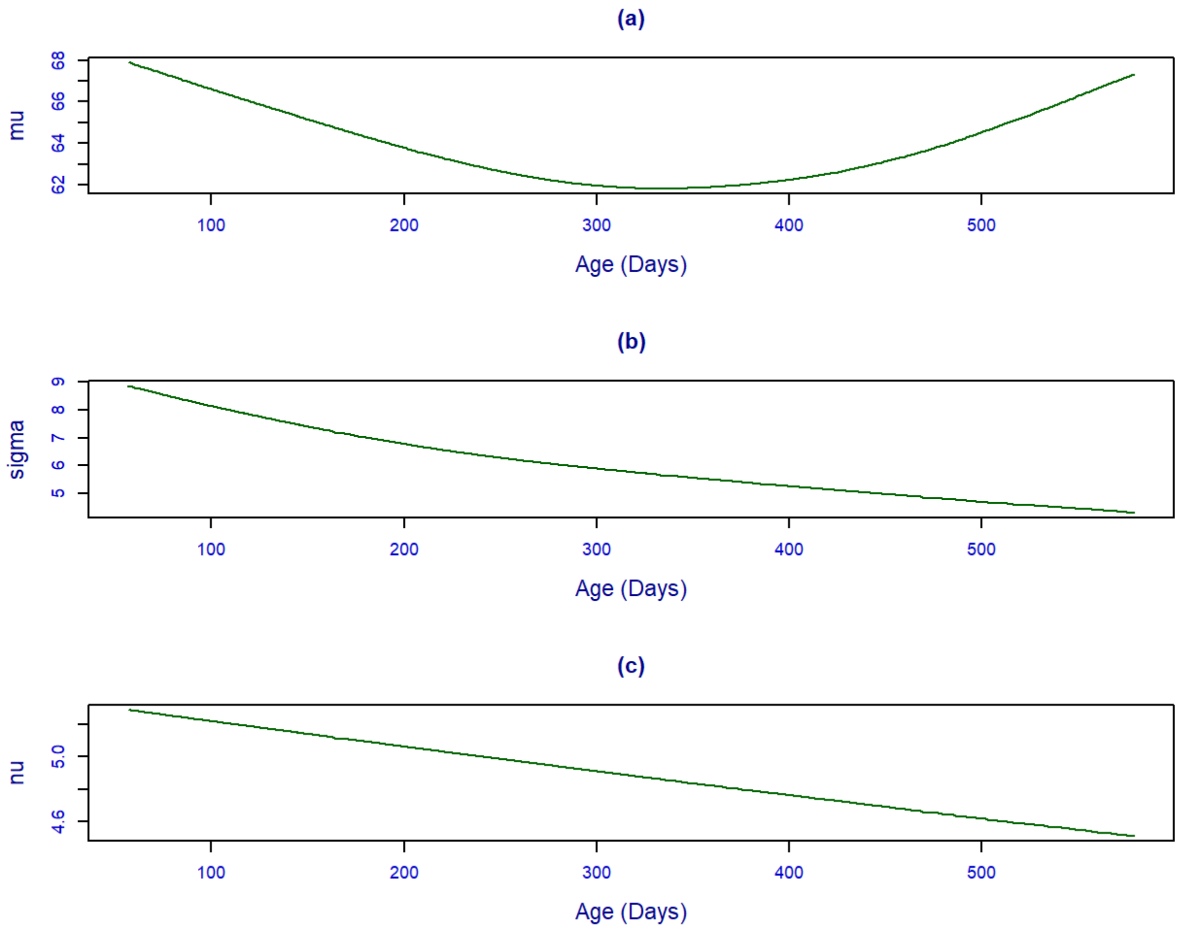

**Figure S17:** N1 Latency fitted term plots of relationship between age and a) mu, b) sigma, and c) nu for the *t*-family growth curve model trained on all datasets.

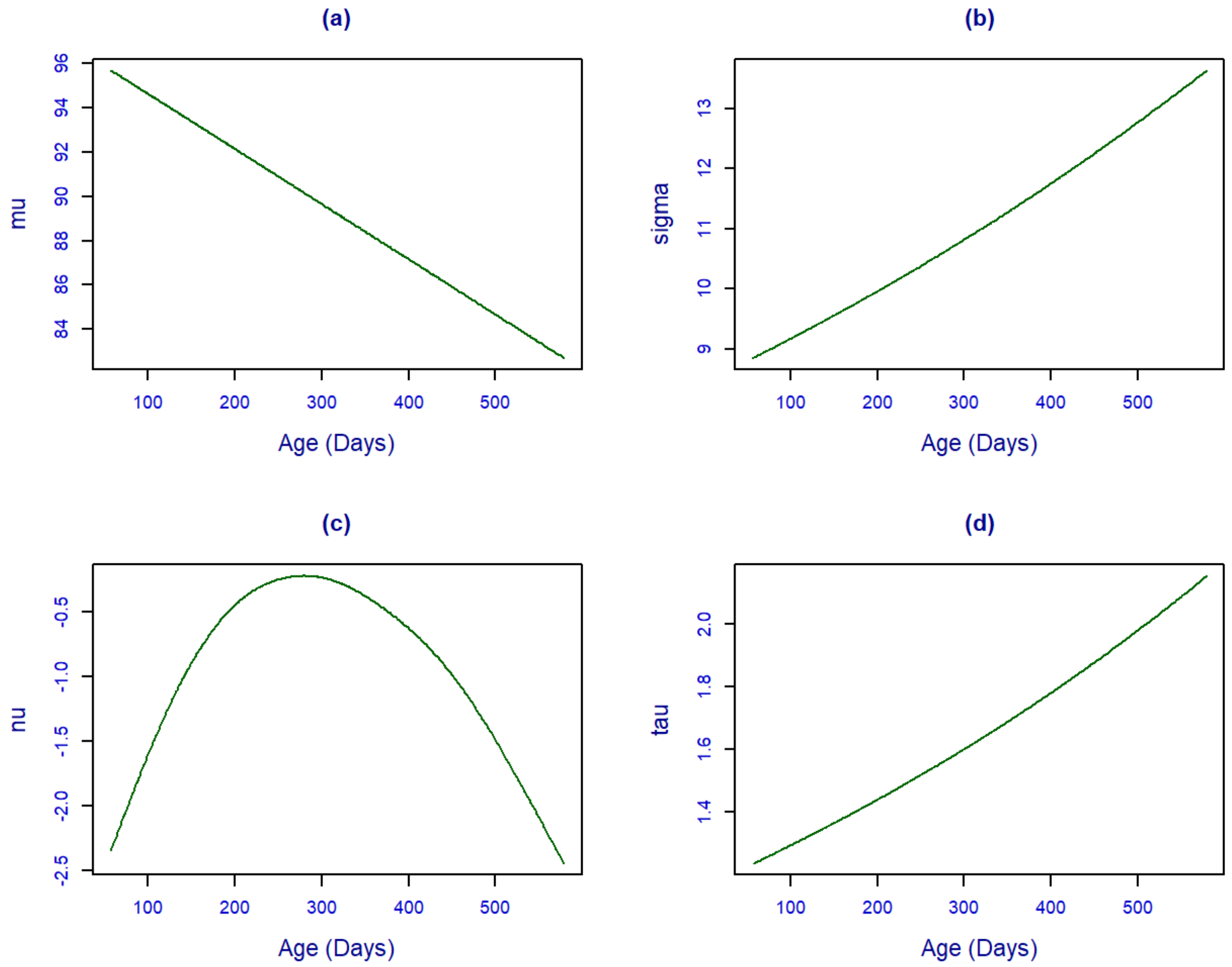

**Figure S18:** P1 Latency fitted term plots of relationship between age and a) mu, b) sigma, c) nu, and d) tau for the Johnson's $S_U$ growth curve model trained on all datasets.

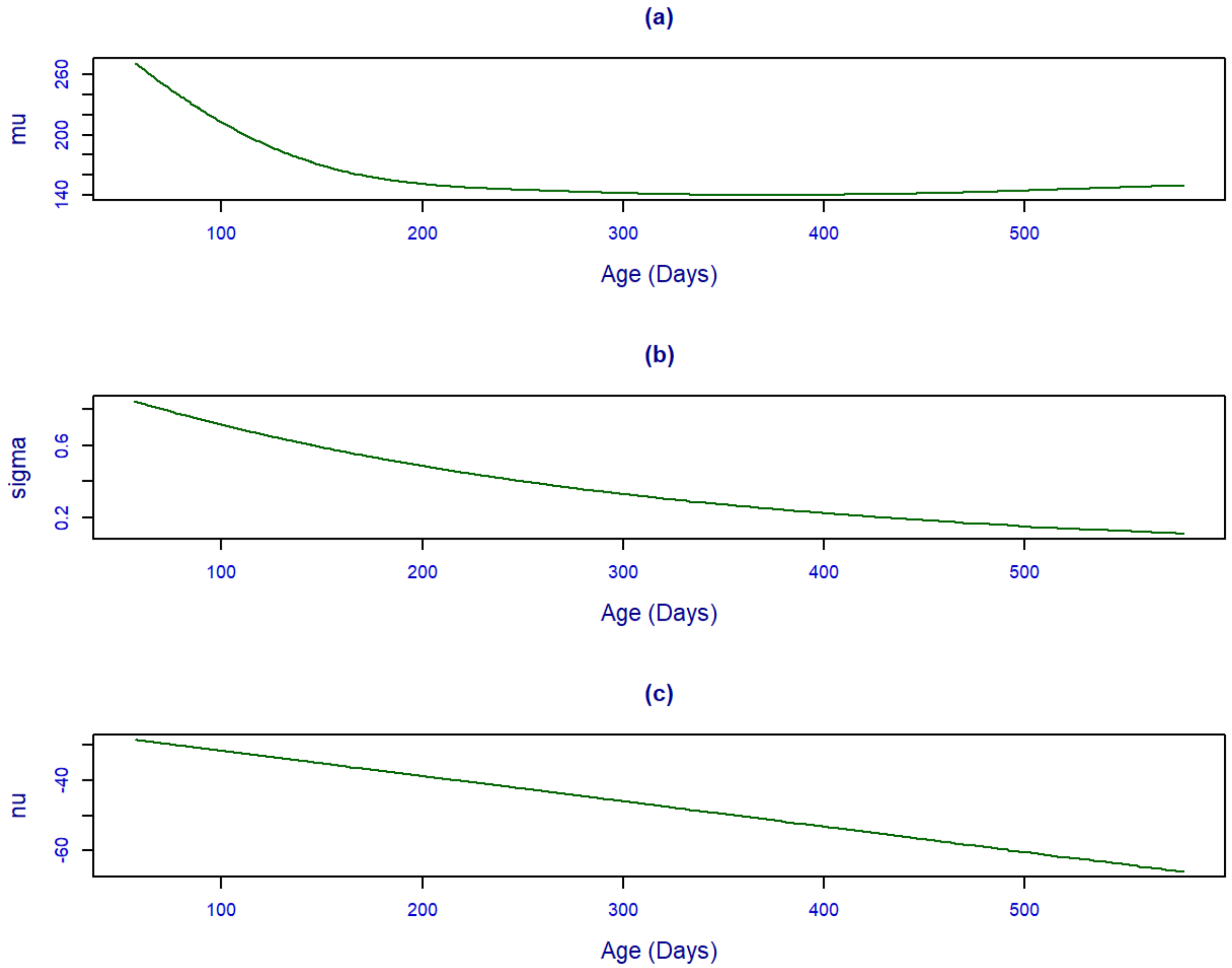

**Figure S19:** N2 Latency fitted term plots of relationship between age and a) mu, b) sigma, and c) nu for the generalized inverse Gaussian growth curve model trained on all datasets.

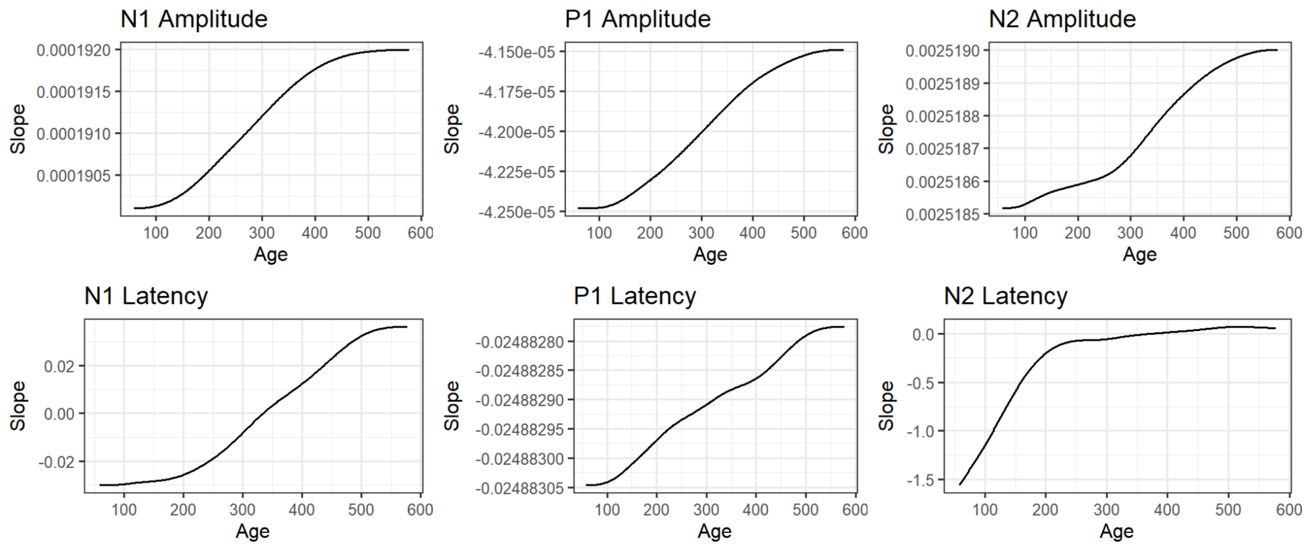

**Figure S20:** Slope plots of the *mu* term estimated from numerical derivation of predicted values from the full models.

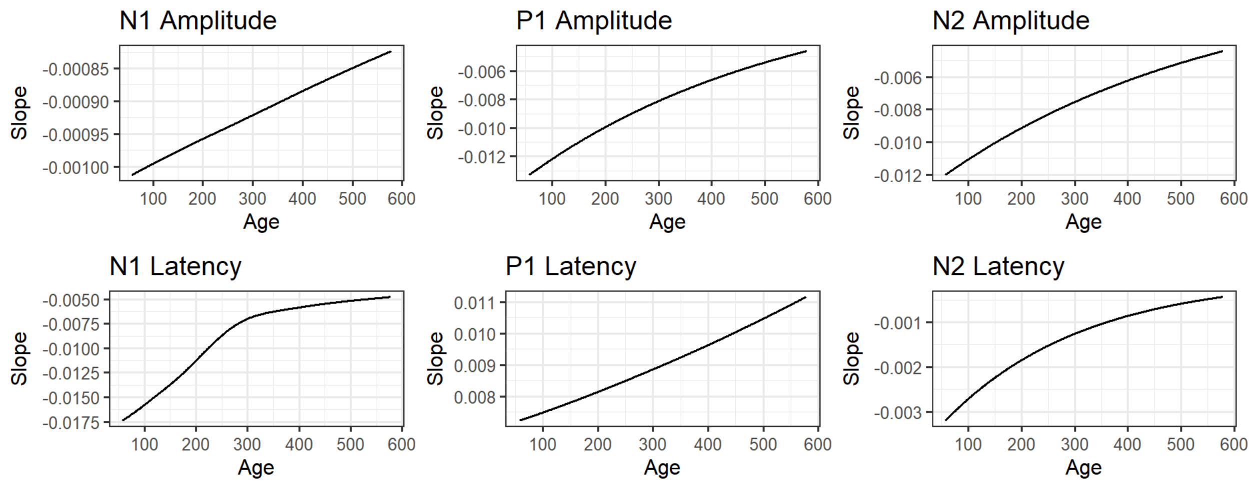

**Figure S21**: Slope plots of the *sigma* term estimated from numerical derivation of predicted values from the full models.

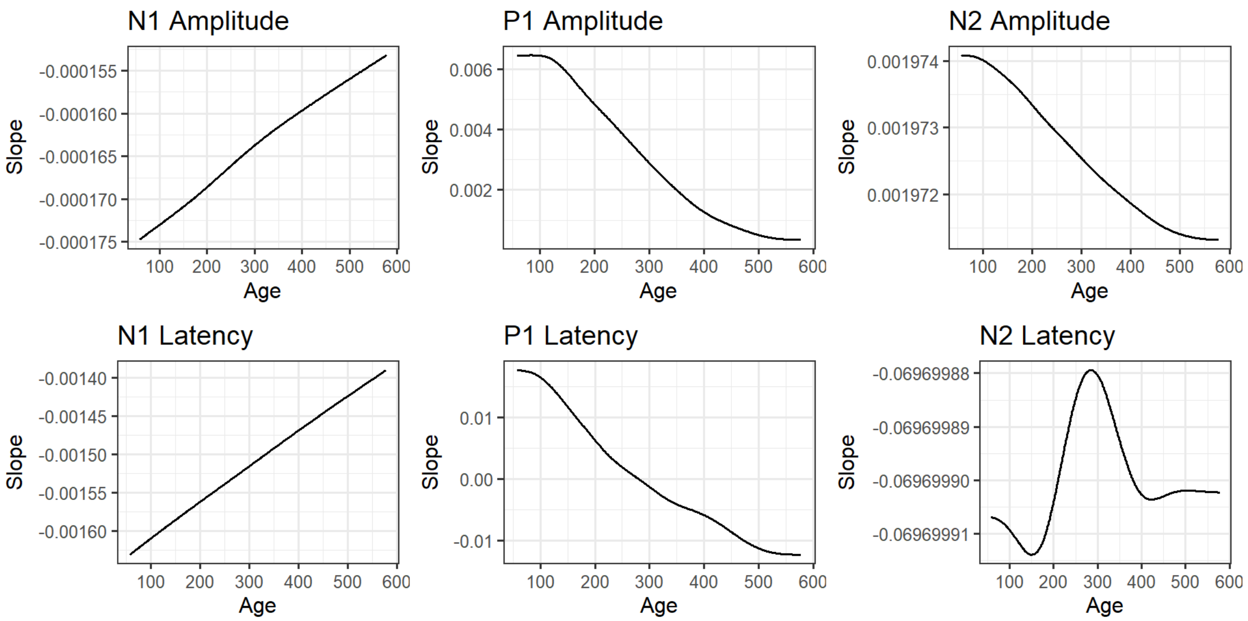

**Figure S22**: Slope plots of the *nu* term estimated from numerical derivation of predicted values from the full models.

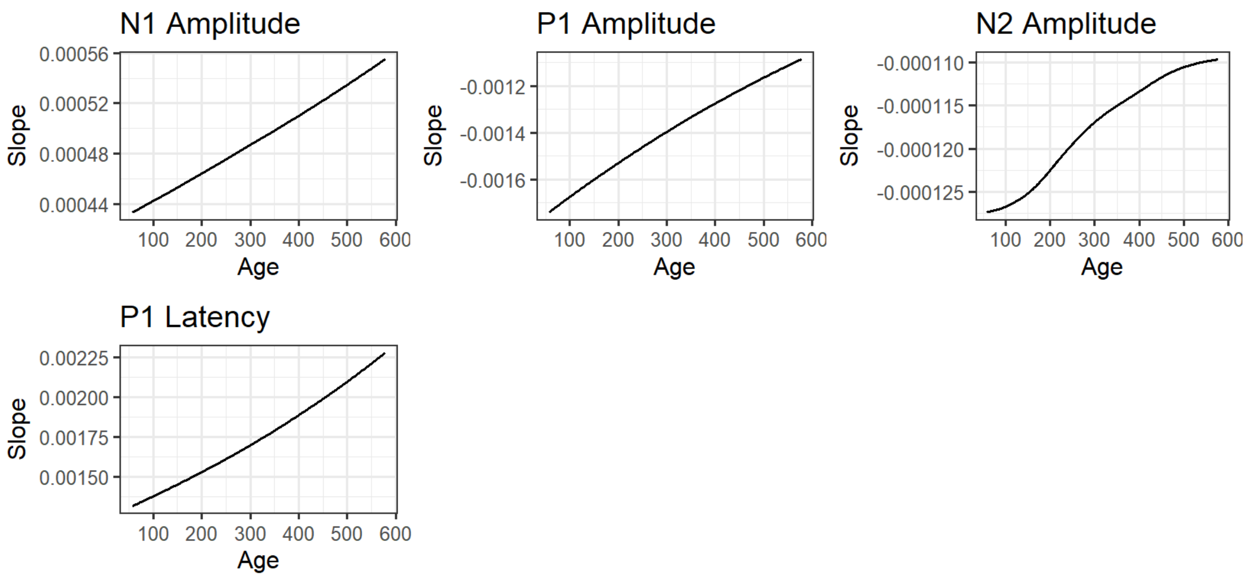

**Figure S23**: Slope plots of the *tau* term estimated from numerical derivation of predicted values from the full models. N1 and N2 Latency models do not have a fourth *tau* parameter.

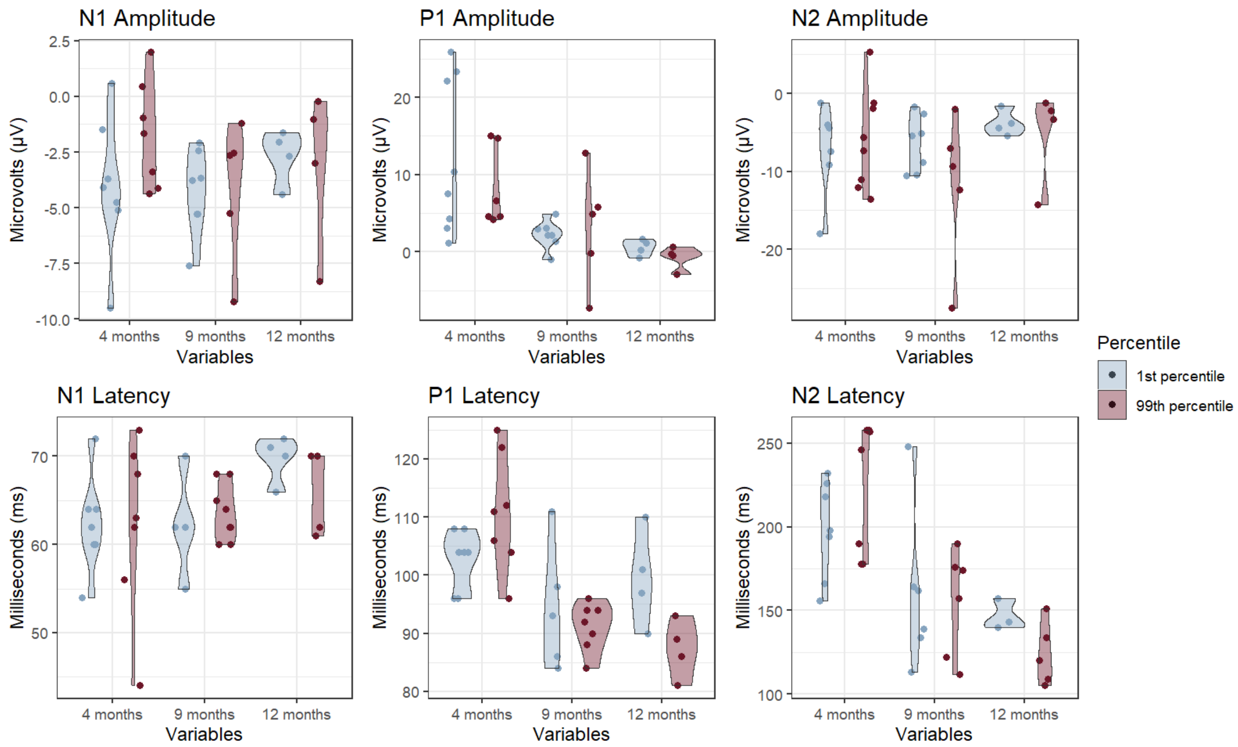

**Figure S24**: Violin plots of VEP feature values comparing 1st and 99th percentile of composite deviation scores at measurement age. Scores estimated using models trained on all three sites. ($n_{4\ months} = 14$; $n_{9\ months} = 12$; $n_{12\ months} = 8$)

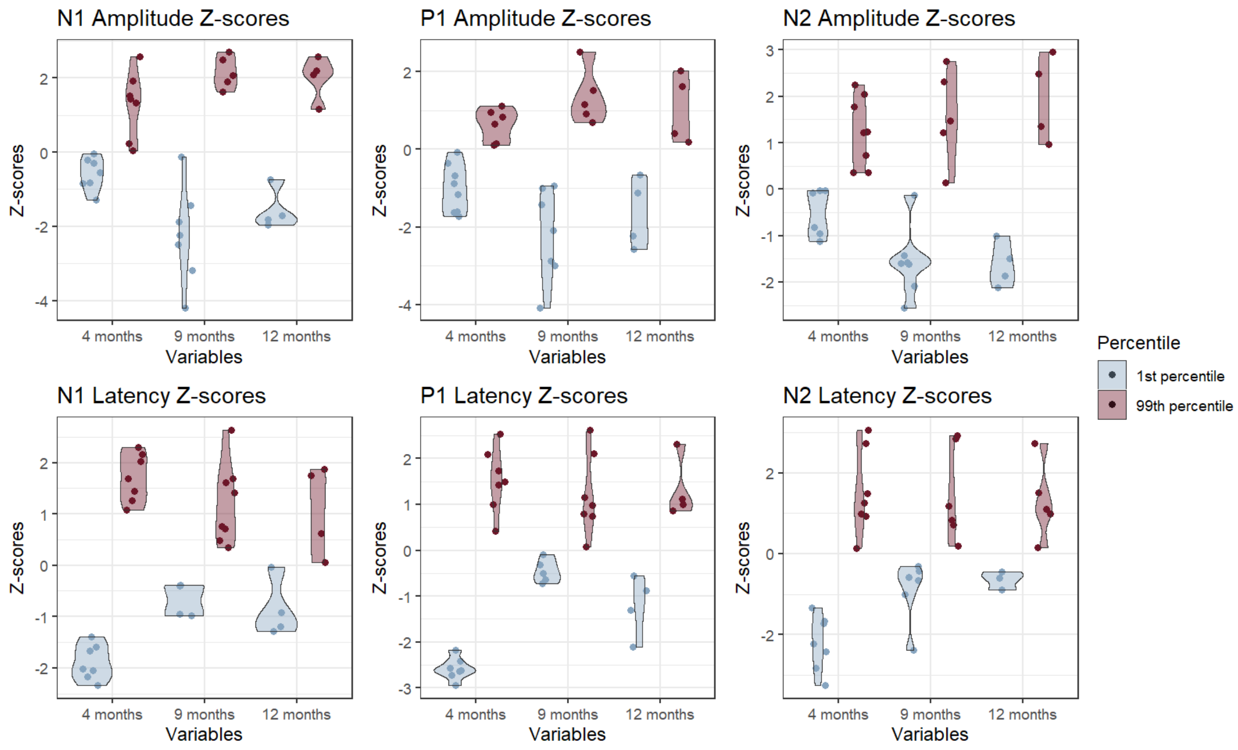

**Figure S25**: Violin plots of VEP feature z-scores comparing 1st and 99th percentile of composite deviation scores at measurement age. Scores estimated using models trained on all three sites. ($n_{4\ months} = 14$; $n_{9\ months} = 12$; $n_{12\ months} = 8$)

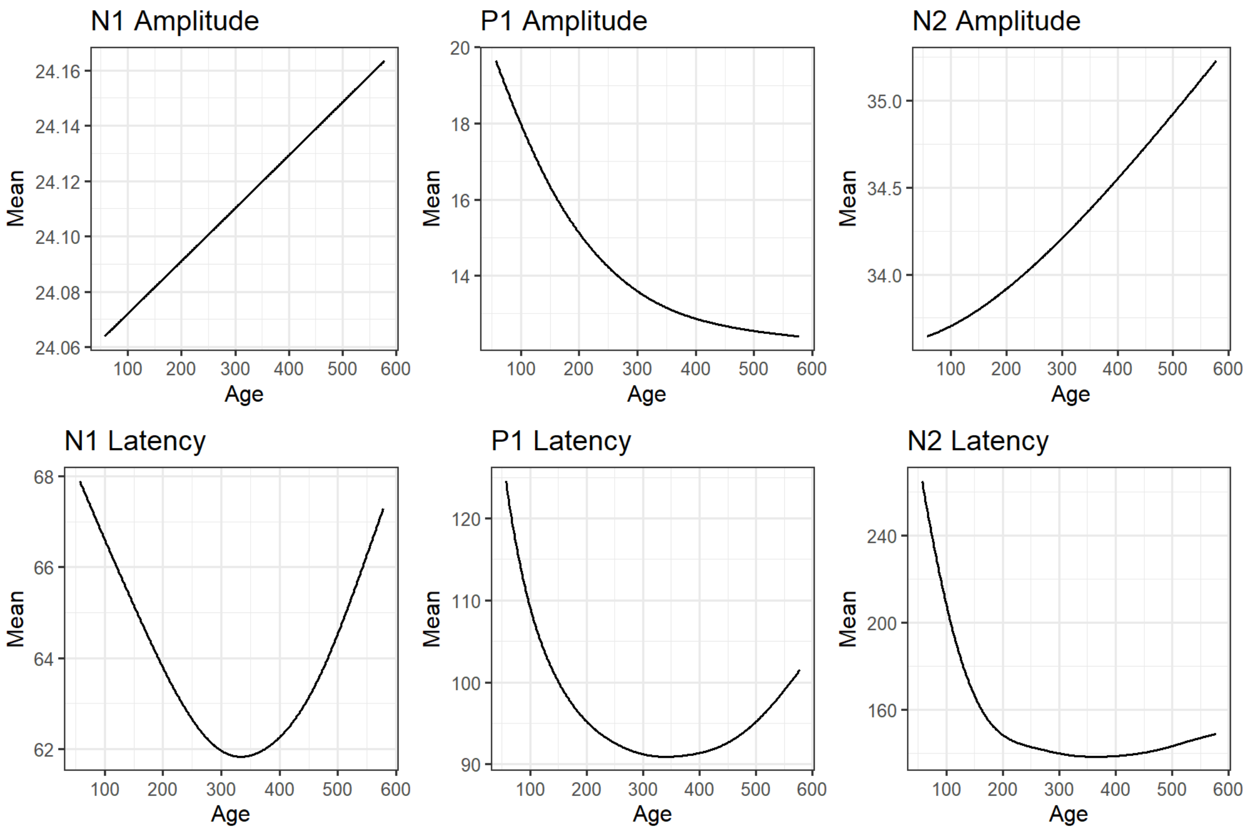

**Figure S26:** Line plots of the 50th percentile of each VEP feature predicted by the full models.

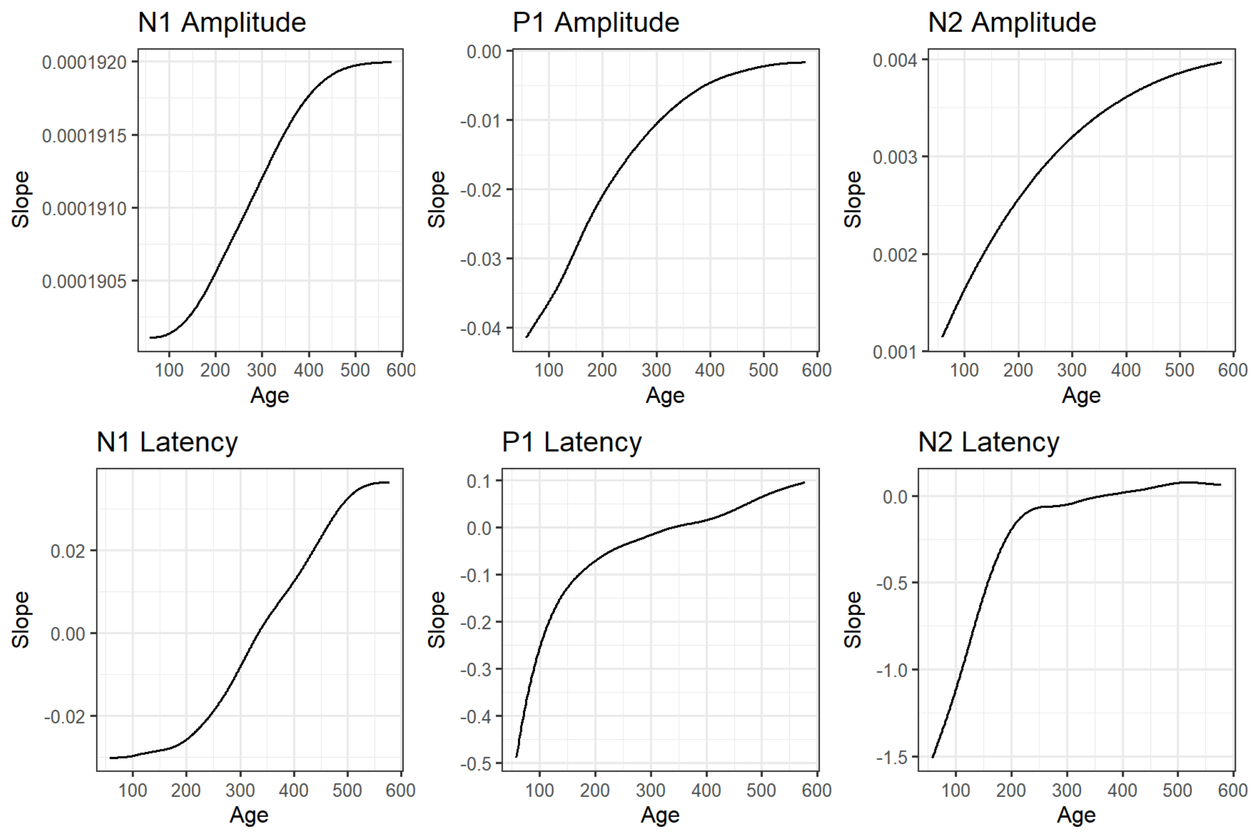

**Figure S27:** Slope plots of the 50th percentile of each VEP feature predicted by the full models.

Slopes were estimated using numerical derivation of the predicted 50th percentile.

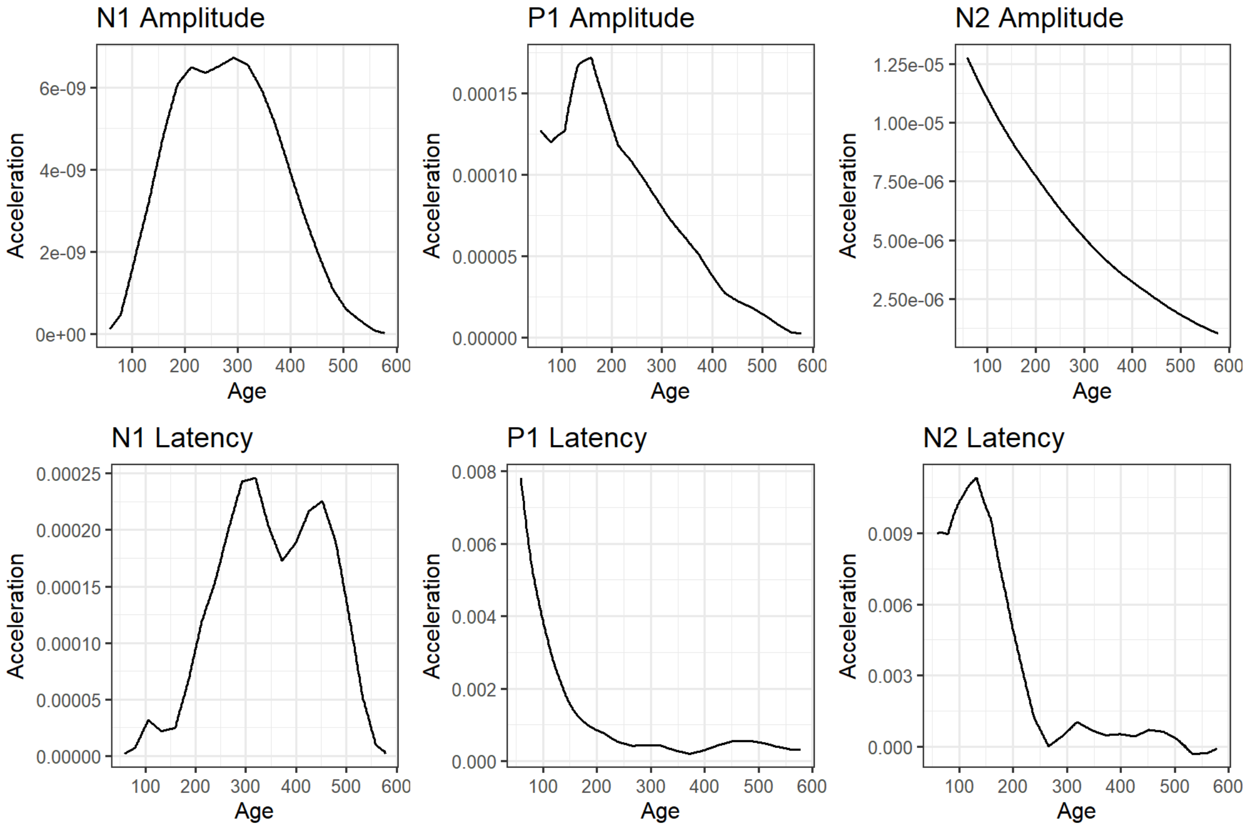

**Figure S28:** Line plots of the second derivative of the 50th percentile of each VEP feature predicted by the full models. Values were calculated using numerical derivation of the first derivative.

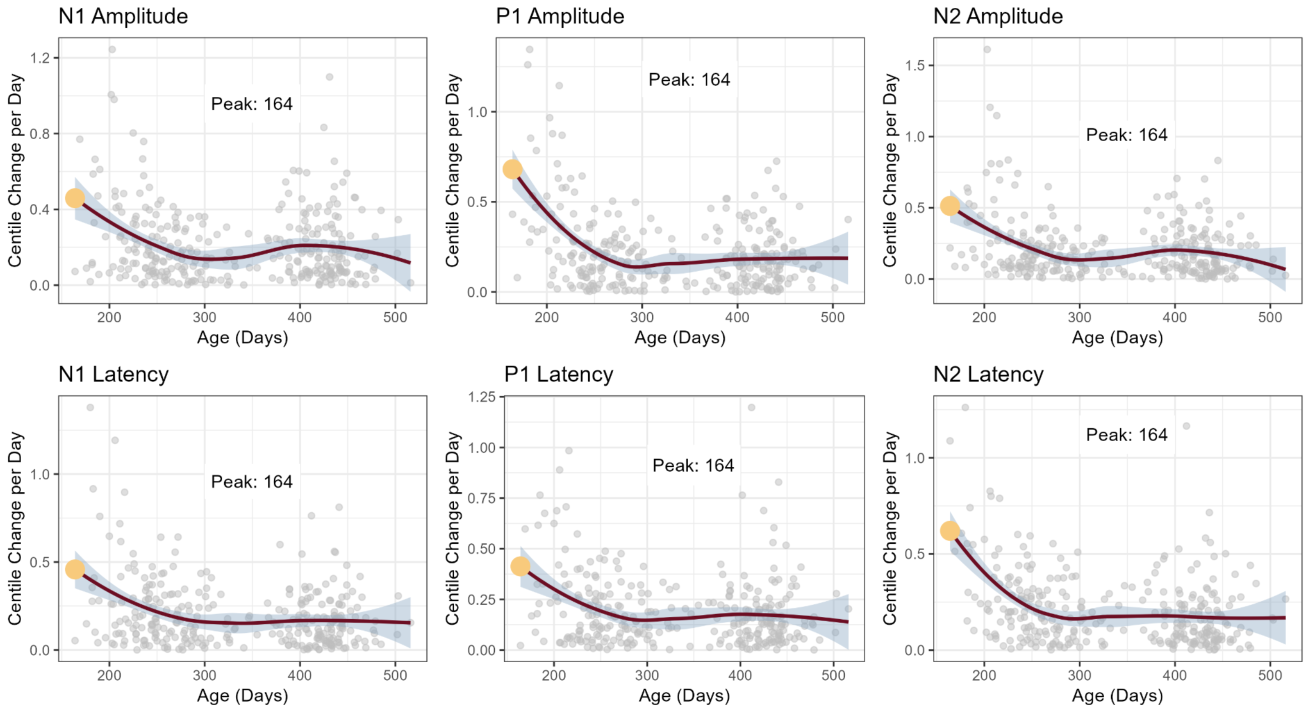

**Figure S29**: LOESS curve (red) fitted to change in centile scores per day for all participants from South Africa site with at least two measurements ($N = 213$).

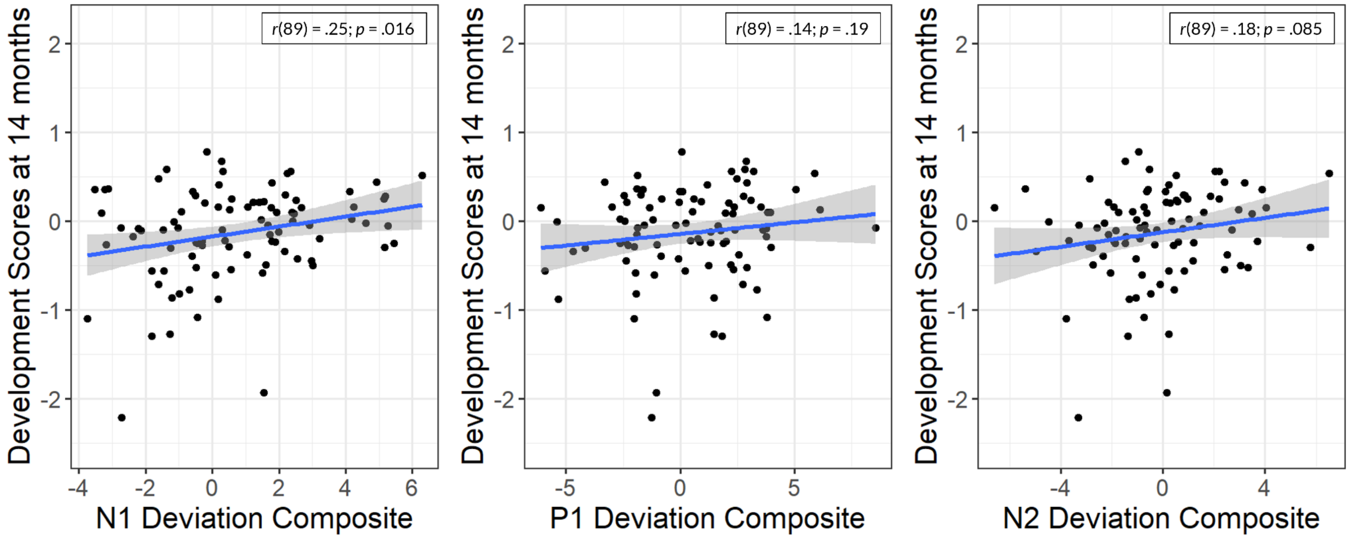

**Figure S30**: Associations between deviations for each feature summed across measurements with Global Scales for Early Development scores at 14 months.

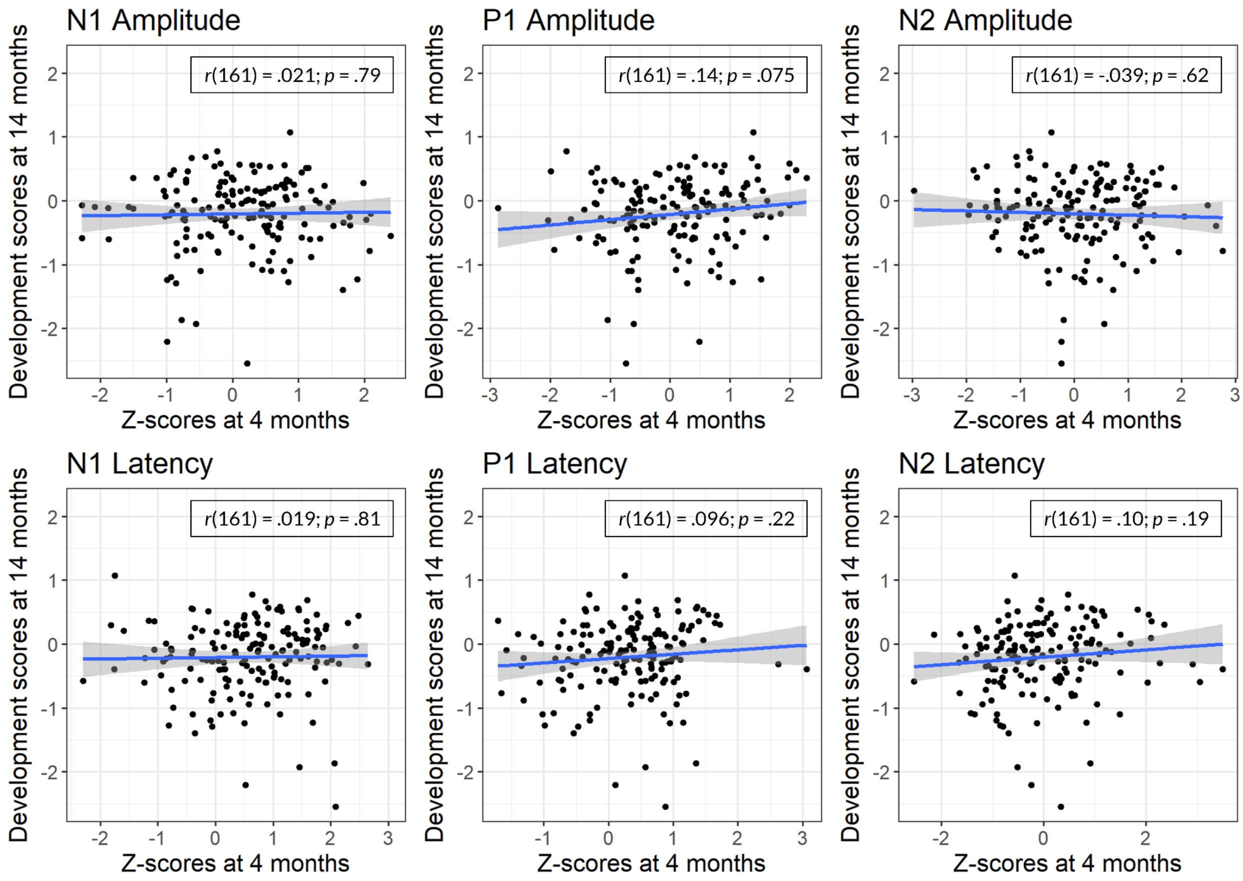

**Figure S31:** Associations between deviations and cognitive outcomes by feature at 4 months

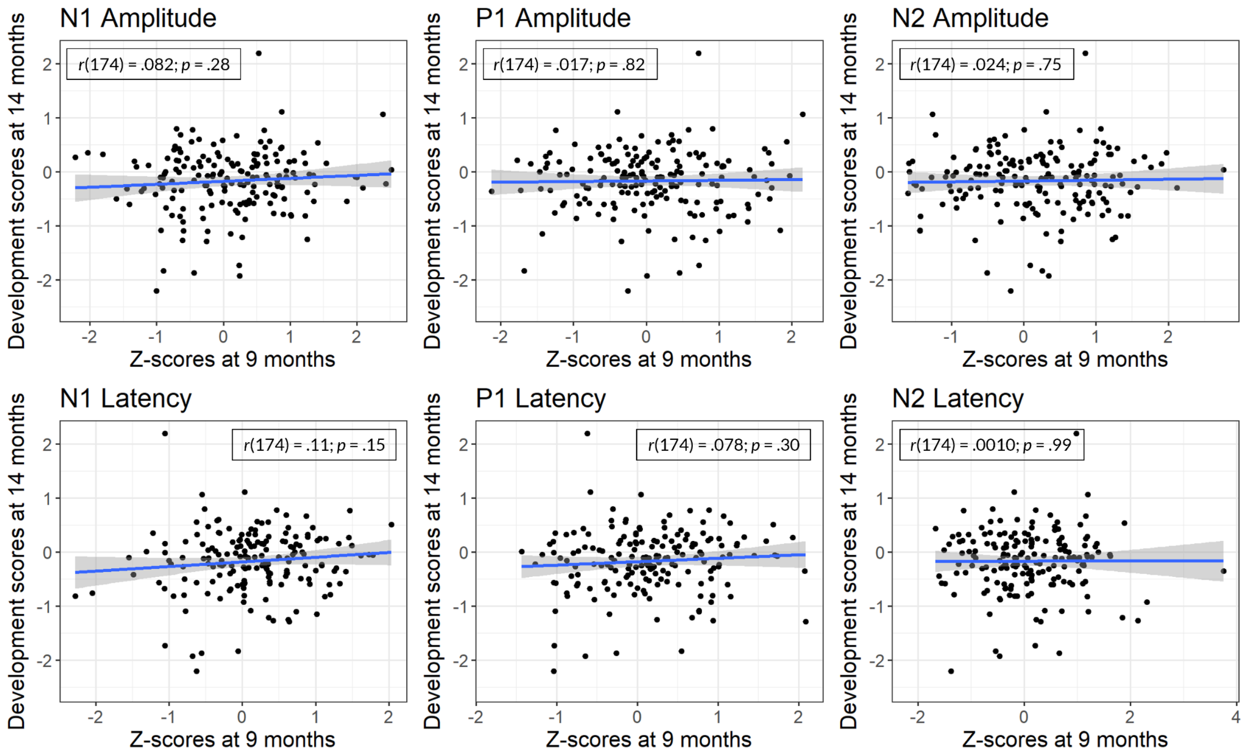

**Figure S32:** Associations between deviations and cognitive outcomes by feature at 9 months

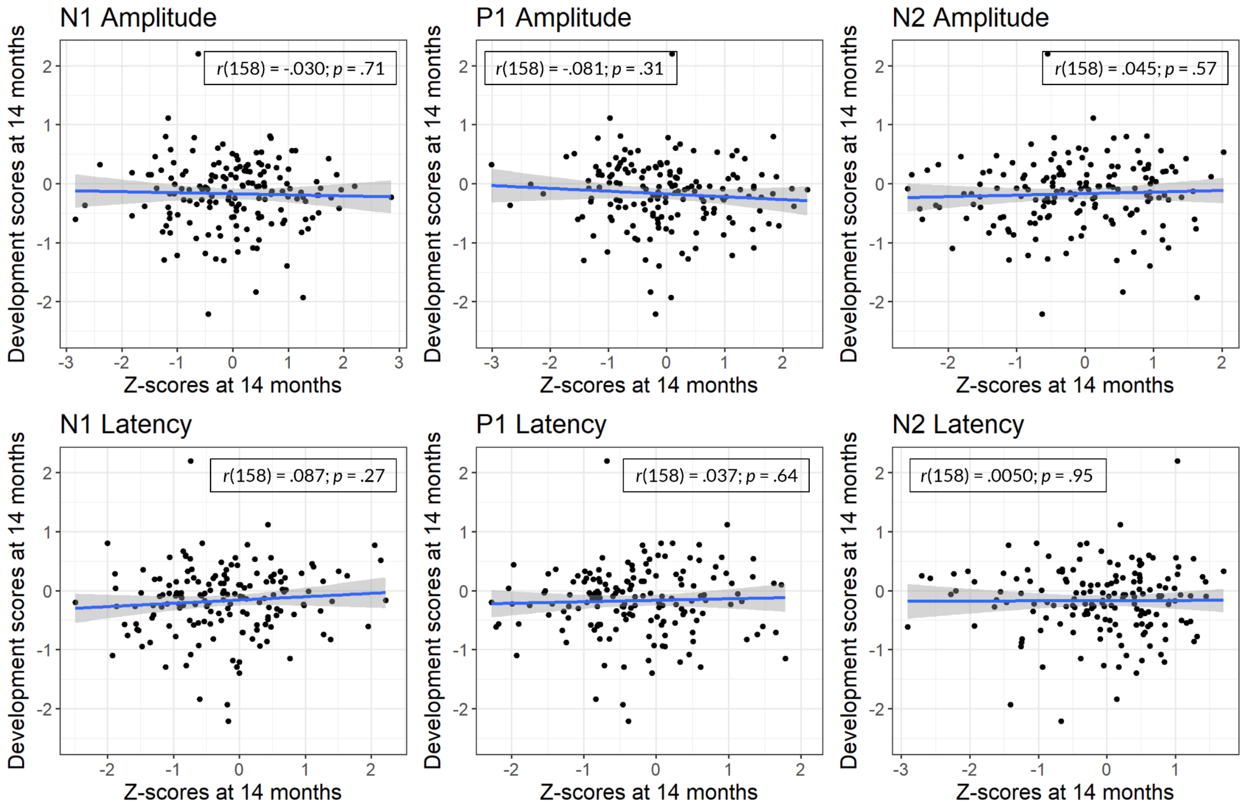

**Figure S33:** Associations between deviations and cognitive outcomes by feature at 14 months

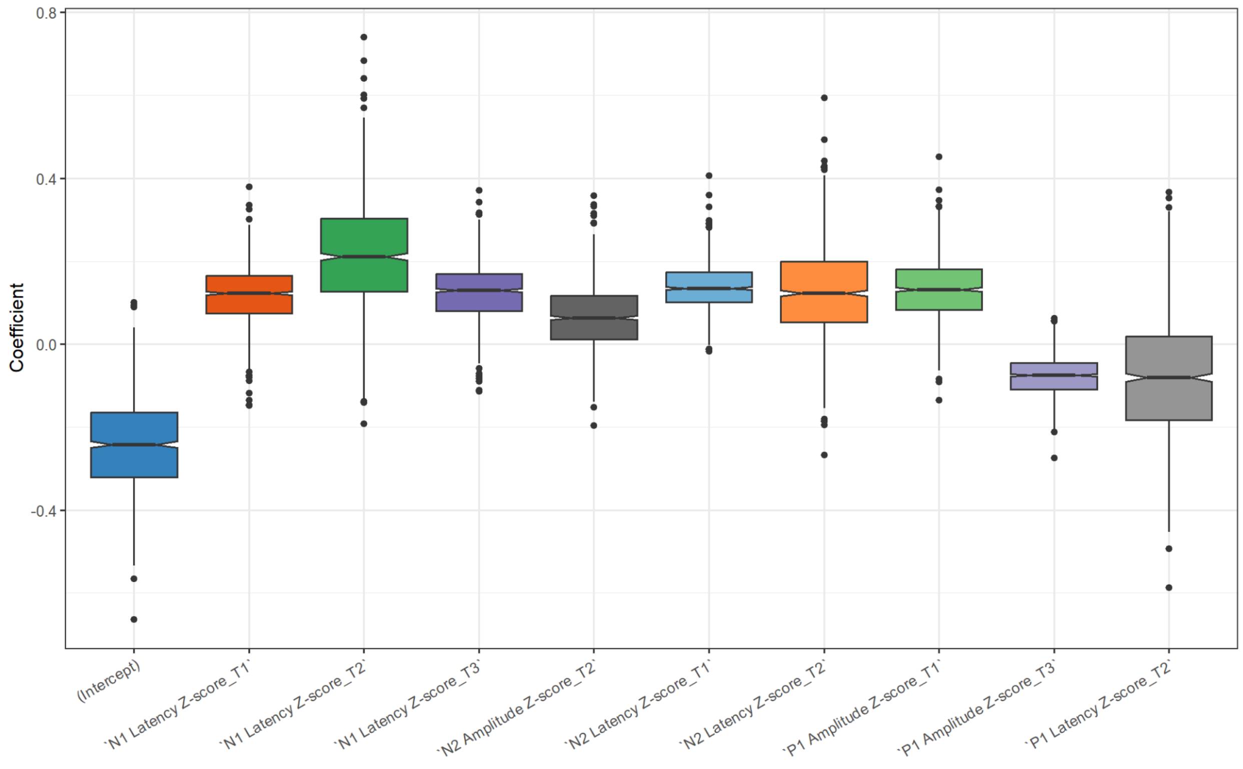

**Figure S34**: Coefficients of linear model predicting GSED scores from VEP feature z-scores. Best terms were selected by maximizing adjusted $R^2$ across an exhaustive search. The best combination of terms was determined through 1000-fold bootstrap with replacement. Coefficient confidence intervals were estimated with 1000-fold bootstrap.
